# Receptor-proximal phosphoform organization precedes β2 integrin activation

**DOI:** 10.64898/2026.11.04.762546

**Authors:** Nicholas A. Hummell, Harathi Jonnagaddala, Marian Kalocsay, Travis I. Moore

## Abstract

Receptor signaling is often framed as protein recruitment after stimulation, but whether receptor-proximal environments are organized in composition and phosphorylation state beforehand remains unclear. We used CD18-APEX2 proximity labeling and a spatial membrane reference to quantify proteins and phosphosites near leukocyte β2 integrins. At rest, receptors, adaptors and Rap GTPases were preferentially represented relative to several catalytic classes. Parent-protein correction revealed spatial phosphoform organization, with 32% of 4,564 phosphosites deviating significantly from the distribution center and 56% of proteins with multiple quantified sites showing divergence beyond measurement error. Sustained activation altered 31% of phosphosites while 95.9% of proteins remained equivalent within ±0.5 log_2_ and spatial phosphoform organization largely persisted. Most phosphorylation responses were shared across both labeling environments, whereas receptor-preferential changes were dominated by phosphosites. These findings suggest that receptor signaling can regulate pre-existing molecular states without extensive protein redistribution.

## INTRODUCTION

Receptor signaling is often studied through the recruitment and assembly of signaling proteins following stimulation. However, the extent to which receptor-proximal environments are organized before stimulation, and whether this organization extends to the phosphorylation states of resident proteins, remains unclear. Leukocyte β2 integrins provide a compelling system in which to examine these questions. Under vascular shear flow, leukocytes can transition from rolling to firm arrest within fractions of a secondsRainger, 1997; Campbell, 1998; Shamri, 2005}. In lymphocytes, lymphocyte function-associated antigen 1 (LFA-1, αLβ2) mediates adhesion through rapid conformational changes that increase ligand-binding affinity without requiring a comparable increase in surface abundance^1–3^. Talin and kindlin-3 regulate these transitions through interactions with the β2 cytoplasmic tail, while Rap1-dependent inside-out signaling coordinates their activity^4–7^. This rapid responsiveness must be balanced against inappropriate adhesion in circulating leukocytes. Whether rapid activation depends on the recruitment of signaling machinery, regulation of proteins already positioned near the receptor, or both remains unresolved.

Biochemical studies have identified talin, kindlin-3 and Rap-associated proteins in association with resting β2 integrins^8,9^, whereas live-cell measurements indicate that talin engagement can be transient or ligand-dependent^10,11^. These findings suggest that signaling proteins may be locally available without continuously engaging the integrin cytoplasmic tail. However, protein proximity alone does not resolve their regulatory state. For example, phosphorylation of the β2 cytoplasmic tail alters its interactions with filamin and 14-3-3 without changing receptor abundance^12^. These observations raise the possibility that rapid integrin regulation involves changes in the phosphorylation states of locally available proteins, in addition to the recruitment of new components.

Distinguishing protein composition from phosphorylation-state organization requires addressing several measurement challenges. Membrane residency contributes to proximity labeling independently of receptor preference, making a spatial membrane reference necessary to distinguish proteins preferentially represented near CD18 from those broadly associated with the cytoplasmic face of the plasma membrane^13,14^. Similarly, phosphosite enrichment may reflect enrichment of its parent protein rather than preferential representation of the phosphorylated form. Correcting phosphosite enrichment for matched parent-protein enrichment separates these contributions and reveals phosphoform representation beyond that predicted by protein proximity^15–17^. Finally, the absence of statistically significant protein changes after stimulation does not establish that protein composition is retained. Equivalence testing provides positive evidence that activation-associated changes remain within a prespecified effect-size interval^18,19^.

We used APEX2 proximity labeling at the cytoplasmic tail of CD18 in Jurkat T cells, together with a spatial membrane reference, to quantify receptor-proximal proteins and phosphosites. Cells were examined at rest on bovine serum albumin (BSA), following engagement with immobilized intercellular adhesion molecule 1 (ICAM-1), or after sustained co-stimulation with ICAM-1 and phorbol 12-myristate 13-acetate (PMA). We asked whether the CD18 environment is organized before activation in both protein composition and phosphorylation state, and whether these organizational features persist following stimulation. Parent-protein correction distinguished spatial phosphoform preferences from protein enrichment, while equivalence testing and comparison with the membrane reference distinguished retained protein architecture from activation-associated remodeling. At rest, the CD18 environment showed preferential representation of selected receptors, adaptors and regulatory proteins, together with spatial phosphoform organization not predicted by parent-protein proximity. Sustained activation extensively remodeled phosphorylation while largely preserving protein composition and spatial phosphoform organization. Most phosphorylation responses were shared across the two labeling environments, whereas the smaller receptor-preferential response was dominated by phosphosites. Together, these findings identify pre-existing protein and phosphoform organization as distinct features of the receptor-proximal signaling environment, suggesting that activation can remodel molecular state without extensive reorganization of local protein composition.

## RESULTS

### A compartment-referenced map of the CD18 cytoplasmic neighborhood

To quantify protein and phosphorylation organization near CD18, we fused APEX2 to the C terminus of CD18 (*ITGB2*) in *ITGB2*-null Jurkat T cells. A separate cell line derived from the same parental background expressed APEX2 bearing the HRAS CAAX membrane-targeting motif, providing a spatial membrane reference. Cells were incubated on BSA, immobilized ICAM-1, or ICAM-1+PMA for 30 min, followed by 1 min of proximity labeling. Three biological replicates consisted of independent cultures labeled on separate days. Tandem mass tag (TMT) mass spectrometry quantified 6,456 proteins and 4,627 phosphosites, with protein and phosphosite measurements derived from the same labeling reactions within a single TMT multiplex (**Fig. 1a**).

**Fig. 1.**
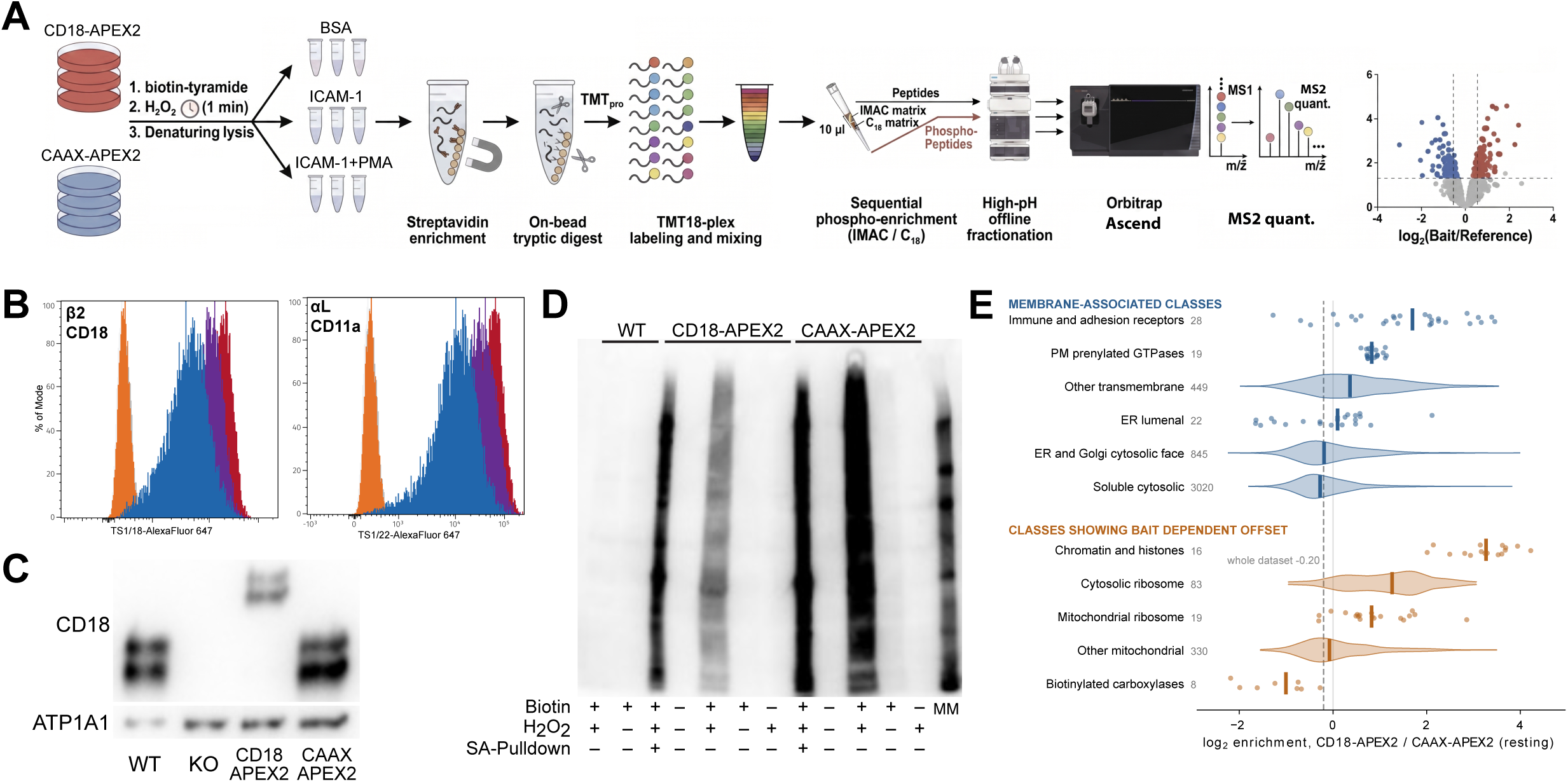
Experimental design and validation of spatially referenced CD18-APEX2 proximity proteomics. **a,** Experimental workflow. APEX2 was fused to the cytoplasmic C terminus of CD18 and compared with membrane-anchored APEX2 bearing the HRAS CAAX targeting motif. Cells were incubated for 30 min under resting BSA, immobilized ICAM-1 alone, or ICAM-1+PMA conditions and labeled for 1 min. Biotinylated proteins were captured using streptavidin, digested on-bead and quantified from TMTpro 18-plex samples by MS2-based Orbitrap Ascend mass spectrometry. Phosphopeptides and the non-phosphopeptide fraction came from the same labeling reactions. **b,** Flow-cytometric surface expression of CD18 (β2; TS1/18) and CD11a (αL; TS1/22) in WT, β2-knockout, CD18-APEX2 and CAAX-APEX2 cells. Histograms are normalized to the mode; gray, secondary antibody only; orange, knockout; red, WT; blue, CD18-APEX2; purple, CAAX-APEX2. Reconstituted cells were sorted to approximate parental expression of both subunits, consistent with restored LFA-1 surface assembly. **c,** Anti-CD18 immunoblot in membrane fractions from WT, β2-knockout (KO), CD18-APEX2 and CAAX-APEX2 lines. The higher-molecular-weight CD18-APEX2 fusion replaces endogenous CD18 in the reconstituted line; CAAX-APEX2 retains endogenous CD18. ATP1A1 serves as the membrane loading control. **d,** Detection of streptavidin-reactive biotinylated proteins in WT, CD18-APEX2 and CAAX-APEX2 samples under the indicated biotin-phenol, H₂O₂ and streptavidin pull-down (SA-Pulldown) conditions. Broad APEX2-dependent labeling required biotin-phenol and H₂O₂, and streptavidin capture enriched biotinylated proteins; MM, molecular weight marker. **e,** Resting BSA log₂ CD18-APEX2/CAAX-APEX2 enrichment by membrane-association class and selected classes showing reference offsets. Each point represents one protein; violins show distributions and numbers give proteins per class. Dashed line, whole-dataset median (−0.20 log₂); solid line, parity.

Surface CD18 and CD11a expression were restored to approximately wild-type (WT) levels (**Fig. 1b**), consistent with LFA-1 assembly and trafficking because surface αL depends on β2 pairing^20^. Membrane fractionation confirmed that CD18-APEX2 replaced endogenous CD18 in the reconstituted line, whereas CAAX-APEX2 cells retained endogenous CD18 (**Fig. 1c**). Biotinylation required APEX2, biotin-phenol and hydrogen peroxide, and streptavidin enrichment recovered the labeled proteins (**Fig. 1d**). Together, these experiments established receptor surface expression, membrane localization and APEX2-dependent labeling.

We use “territory” to describe the proteins sampled within the APEX2 labeling range without implying a defined physical boundary or stable membership in a molecular complex. Because proximity labeling integrates encounters over time, proteins detected near the CD18 cytoplasmic tail may be directly associated with the receptor or transiently present within the labeling volume^21,22^. Within each condition, log_2_(CD18/CAAX) enrichment measures relative protein representation near CD18 compared with the spatial membrane reference. Positive values indicate preferential representation in the CD18 sample, whereas negative values indicate greater representation in the CAAX reference; neither establishes direct receptor binding. Protein abundance percentile instead reflects relative abundance within a single proximity-labeled sample. Interpretation of these measurements must also account for systematic differences between the separately generated cell lines. Endogenously biotinylated carboxylases, which are captured independently of APEX2 labeling and should therefore show no CD18/CAAX difference, were offset toward CAAX. Enrichment also varied with membrane-association class: prenylated plasma-membrane GTPases were shifted toward CD18 and early secretory-pathway proteins toward CAAX (Fig. 1e; Supplementary Fig. 1). Enrichment was calculated identically for every protein, but these class-level offsets mean that values are most informative when proteins are compared with others in the same membrane-association class rather than read as absolute measures of receptor association.

### The resting β2 integrin territory already contains a structured protein architecture

At rest, CD18 sampled a compositionally structured membrane environment relative to the CAAX reference (**Figs. 2a,b**). Receptors and docking adaptors preferentially represented near CD18 included PTPRCAP (+3.45), LAIR1 (+3.39), SLAMF6 (+2.25), SIT1 (+2.11), LAT (+2.05), PAG1 (+1.62), TRAT1 (+1.59) and CD84 (+1.29), all with Benjamini–Hochberg-adjusted q ≤ 8.8 × 10⁻⁸ (**Figs. 2c,d**). L-selectin (+3.17), ICAM-2 (+2.86), CD43 (+2.68) and four tetraspanins were also enriched. Immune and adhesion receptors showed greater enrichment than other transmembrane proteins (median +1.70 versus +0.37; n = 28 and 449; P = 7.2 × 10⁻⁶; **Fig. 1e**). All six previously described β2 integrin-associated surface proteins quantified in this study (L-selectin, CD47, ICAM-2, CD43, CD147 and ICAM-3) were enriched relative to CAAX^9^. CD47, previously identified in cis proximity to β2 integrins^23^, showed enrichment of +3.10 across 15 peptides, whereas the abundant transferrin receptor showed little preference (−0.30; **Fig. 2b**). These measurements reflect proximity to cytoplasmic receptor domains rather than direct molecular interactions.

**Fig. 2.**
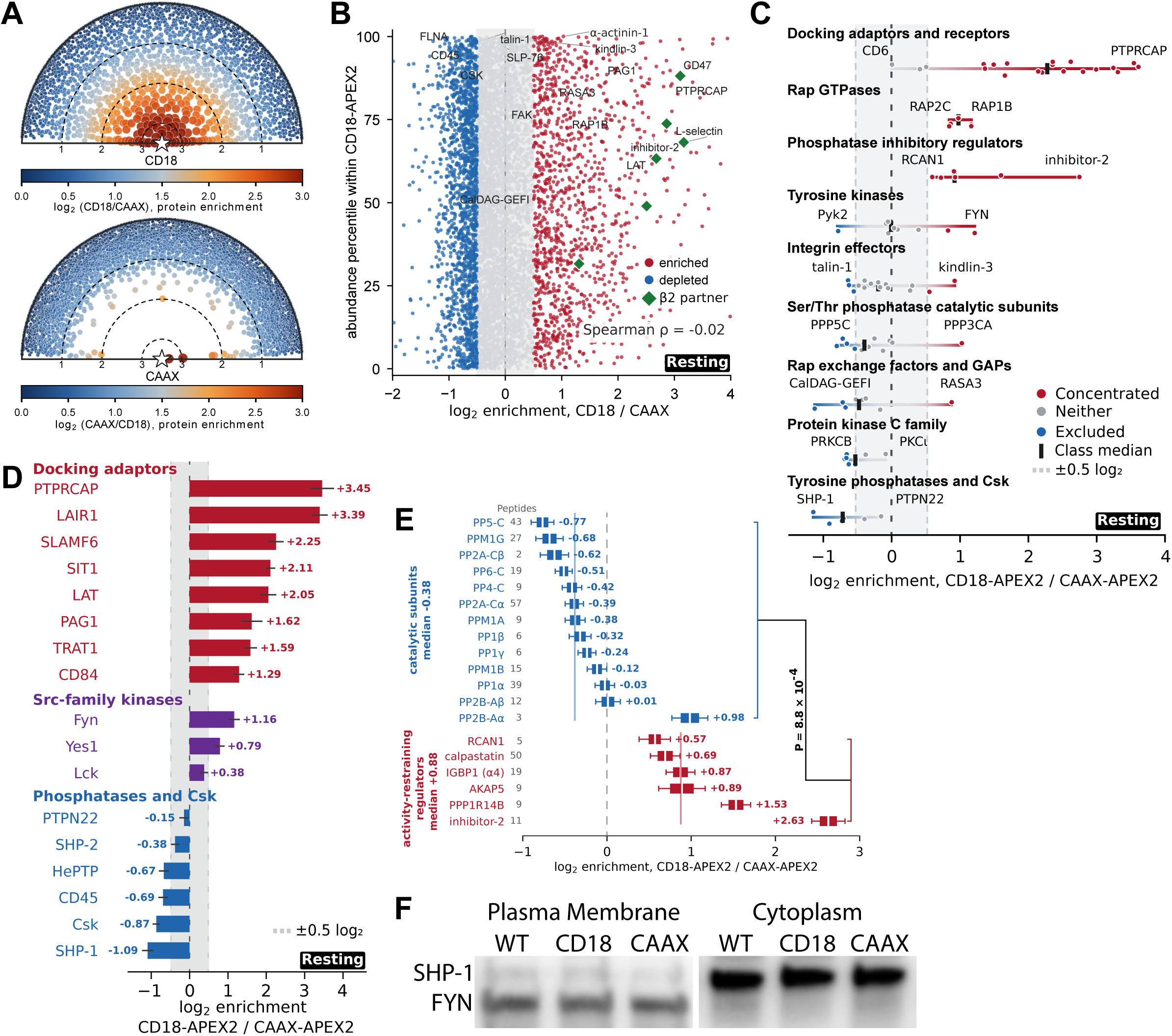
The resting CD18 territory contains a structured protein architecture. **a,** Radial visualization of resting protein enrichment in three biological replicates for CD18-APEX2 and CAAX-APEX2. Color indicates log₂ enrichment of the specified sample relative to the other labeling sample; the CD18 plot uses CD18/CAAX, while the CAAX plot uses CAAX/CD18. Distance from the central bait symbol decreases with enrichment (dashed rings at 1, 2 and 3 log₂; proteins with enrichment ≤ 0 lie at the rim), and angle carries no information. **b,** Protein-abundance percentile within the CD18 sample plotted against resting log₂ CD18/CAAX enrichment across 6,456 quantified proteins. Red and blue show features passing q < 0.05 and |log₂ enrichment| ≥ 0.5 in the respective directions; green diamonds indicate the six previously reported β2 integrin-associated surface partners. Spearman ρ = −0.02. **c,** Resting enrichment of all quantified members of nine curated signaling classes; the protein at each end of a class is labeled. Red, gray and blue denote proteins concentrated at CD18, neither, or excluded (q < 0.05 and |log₂ enrichment| ≥ 0.5); black marks show class medians and the shaded band marks ±0.5 log₂. **d,** Resting enrichment of the docking adaptors, Src-family kinases, tyrosine phosphatases and Csk discussed in the Results. Bars show fitted log₂ enrichment with model-based uncertainty; colors denote these groups and the shaded band marks ±0.5 log₂. **e,** Resting enrichment of 13 PPP- and PPM-family Ser/Thr phosphatase catalytic subunits (blue) versus six selected activity-restraining and regulatory proteins (red). Numbers indicate quantified peptides; median enrichment −0.38 and +0.88 log₂, respectively (two-sided Mann–Whitney P = 8.8 × 10⁻⁴). **f,** Immunoblots detecting SHP-1 and FYN in resting plasma-membrane and cytoplasmic fractions from WT, CD18-APEX2 and CAAX-APEX2 cells. Their different proximity enrichments (FYN +1.16; SHP-1 −1.09 log₂) coexist with similar gross fractionation profiles. Proteomic data derive from three biological replicates (independent cultures labeled on separate days).

Protein abundance within the CD18 sample was nearly independent of CD18/CAAX enrichment (Spearman ρ = −0.02 across 6,456 proteins; **Fig. 2b**). In contrast, ranking abundance within the CAAX reference produced an apparent negative relationship (ρ = −0.38; **Supplementary Fig. 2a**), as expected because CAAX abundance is also the denominator of the enrichment ratio. Talin-1 illustrates the distinction between local abundance and receptor-preferential representation. Although talin ranked at the 97.8th abundance percentile within CD18-labeled material, it was relatively under-represented compared with CAAX (−0.61, q = 1.5 × 10⁻⁵). Other proteins associated with mature mechanotransduction were similarly detected without preferential enrichment, including Pyk2 (−0.76), filamin A (−0.59), vinculin (−0.53), paxillin (−0.38), LIMS1 (−0.38) and ILK (−0.19). Focal adhesion kinase and zyxin were near parity. By contrast, α-actinin-1 (+1.03) and kindlin-3 (+0.88) were both abundant and enriched, whereas CD84 and TRAT1 were enriched despite ranking near the 30th abundance percentile. Thus, protein abundance within the CD18 environment did not necessarily predict receptor-preferential representation. Protein and phosphosite enrichment landscapes at rest and after ICAM-1+PMA are shown in **Supplementary Figs. 3a–d**.

The resting territory also exhibited functional asymmetry among signaling regulators (**Fig. 2c**). All five quantified Rap isoforms were enriched near CD18 (RAP1B and RAP2B, +1.12; RAP2C, +0.81; q < 3.5 × 10⁻⁶), whereas no quantified Rap exchange factor showed comparable enrichment, including CalDAG-GEFI (−1.07). Most Rap-directed GTPase-activating proteins (GAPs) were under-represented, with RASA3 (+0.84) as an exception. A similar separation was observed between enriched docking adaptors and several reported regulatory enzymes^24–27^. Csk (−0.87), SHP-1 (−1.09), CD45 (−0.69), HePTP (−0.67) and SHP-2 (−0.38) were under-represented, whereas FYN (+1.16) and YES1 (+0.79) were enriched, indicating that this pattern did not extend uniformly across kinase classes. Class-level analyses supported these differences (**Supplementary Fig. 4a–c**). Thirteen selected PPP/PPM Ser/Thr phosphatase catalytic subunits showed a median enrichment of −0.38, compared with +0.88 for six activity-restraining regulators from the PP1, PP2A, calcineurin and calpain systems (P = 8.8 × 10⁻⁴; **Fig. 2e**).

These functional differences must be interpreted in the context of membrane-association and cell-line reference offsets. Although plasma-membrane proteins generally showed greater representation in the CD18 sample, the transmembrane phosphatase CD45 was relatively under-represented (−0.69). This establishes a relative difference between the two proximity environments but does not demonstrate selective exclusion after accounting for membrane association or peptide coverage (**Supplementary Fig. 4b**). Moreover, APEX2 integrates proximity over time rather than measuring stable molecular occupancy. Principal-component analysis of the complete 18-channel protein profiles independently recovered the same broad functional ordering without using class membership to construct the axes (permutation P ≤ 1.5 × 10⁻⁴; **Supplementary Figs. 5a,b**). Immunoblotting showed comparable SHP-1 and FYN abundance in plasma-membrane and cytoplasmic fractions from WT, CD18-APEX2 and CAAX-APEX2 cells (**Fig. 2f**), indicating that their opposite proximity enrichments were not readily explained by gross differences in membrane partitioning between cell lines.

Together, these results establish a compositionally structured resting CD18 territory in which receptors, docking adaptors, Rap GTPases and selected regulatory proteins are preferentially represented relative to several catalytic enzyme classes.

### Parent-protein correction reveals spatial phosphoform organization at rest

We next asked whether the phosphorylated forms of proteins near resting CD18 exhibit spatial preferences beyond those predicted by parent-protein proximity. Because phosphosite enrichment can reflect enrichment of its parent protein, we corrected each phosphosite for its matched parent-protein enrichment measured from the same labeling reactions (**Figs. 3a,b**). For 4,564 phosphosite– parent pairs representing 2,015 gene products, we defined the phosphospatial index (PSI) as the log_2_ CD18/CAAX phosphosite ratio minus the corresponding log_2_ parent-protein ratio within each condition^15–17^. Positive PSI indicates greater phosphoform representation near CD18 than predicted by parent-protein proximity. Because PSI compares two proximity-labeled populations, it measures relative phosphoform representation rather than phosphorylation occupancy per molecule. Spatial offsets shared by a phosphosite and its parent protein are removed by the subtraction.

**Fig. 3.**
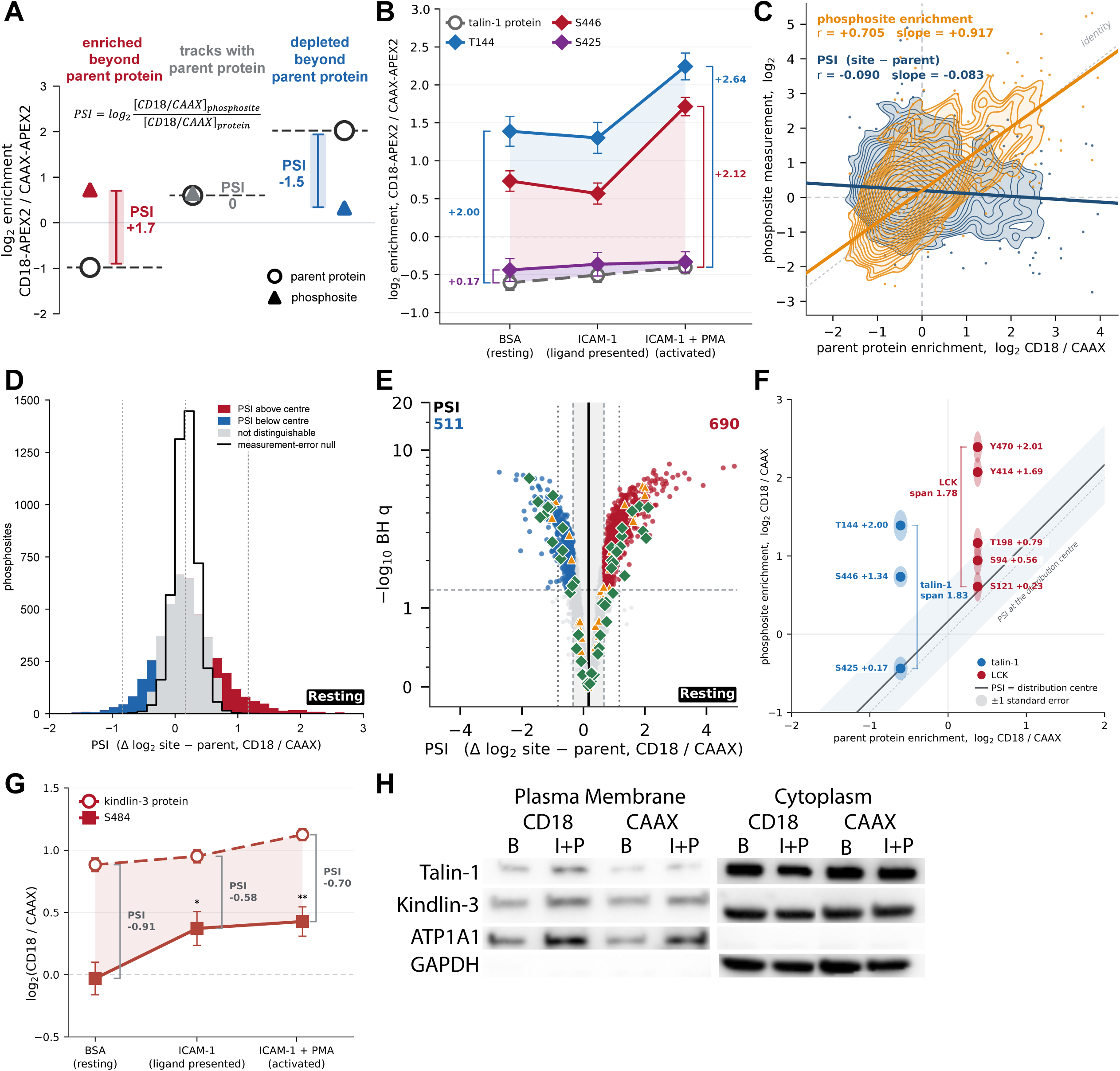
Parent correction reveals spatially selected phosphoforms in the resting CD18 territory. **a,** Definition of the phosphospatial index (PSI) as log₂ CD18/CAAX enrichment of a phosphosite minus the log₂ enrichment of its parent protein. PSI describes relative selection of a phosphoform beyond parent-protein proximity and is not a phosphorylation-occupancy measurement. **b,** Enrichment of talin-1 protein and its T144, S446 and S425 phosphopeptides across BSA, ICAM-1 and ICAM-1+PMA; brackets denote PSI. Points show fitted estimates and model-based uncertainty. **c,** Raw phosphosite enrichment (orange) and PSI (blue) plotted against matched parent enrichment; density contours and regression lines show the corresponding relations. **d,** Distribution of resting PSI relative to a measurement-error null; red and blue distinguish sites above and below the parent-corrected distribution center, while gray indicates unresolved sites; dotted lines mark the center and ±1 log₂ from it. **e,** Significance of resting PSI deviations from the empirical center, with vertical lines marking the center, ±0.5 and ±1 log₂ from it. In all, 690 sites lay above and 511 below the center at q < 0.05 and at least 0.5 log₂ from it. Green diamonds and orange triangles mark sites on immune and adhesion receptors and on the curated adhesion-signaling set; the vertical axis is nonlinear above 1. **f,** Within-protein PSI variation on talin-1 and Lck. The diagonal indicates the empirical center and the shaded band propagated uncertainty; labels give PSI and brackets show within-protein spread. **g,** Kindlin-3 protein enrichment and S484 phosphoform enrichment across conditions. PSI remained negative despite enrichment of the parent protein. Asterisks mark conditions in which S484 enrichment differed from parity (*q < 0.05; **q < 0.01). **h,** Talin-1 and kindlin-3 immunoblots in plasma-membrane and cytoplasmic fractions of CD18-APEX2 and CAAX-APEX2 cells after resting BSA (B) and ICAM-1+PMA (I+P). ATP1A1 and GAPDH serve as fractionation controls. PSI was calculated for 4,564 site-parent pairs representing 2,015 parent gene products.

Raw phosphosite enrichment closely tracked parent-protein enrichment (r = +0.705, slope +0.917), whereas PSI showed little remaining association with parent enrichment (r = −0.090, slope −0.083; **Fig. 3c**). Because protein and phosphosite measurements were normalized independently, PSI was interpreted relative to its empirical distribution center (+0.167 log_2_ at rest) rather than zero (**Supplementary Fig. 6a**). A sensitivity analysis using protein-anchored normalization yielded consistent spatial conclusions (**Supplementary Figs. 7a,b**).

The resting CD18 territory exhibited substantial phosphoform heterogeneity beyond measurement uncertainty. Of 4,564 phosphosite–parent pairs, 1,462 (32.0%) deviated significantly from the empirical PSI center (q < 0.05), and 344 differed by more than 1.0 log_2_, compared with 0.06% expected from fitted measurement errors (**Figs. 3d,e**). Approximately 90% of the observed resting PSI variance remained unexplained by propagated standard errors. The primary analysis conservatively treated phosphosite and parent-protein errors as independent. Direct modeling of per-channel phosphosite-minus-parent differences, accounting for their covariance, produced nearly identical PSI estimates (r = 0.998), smaller standard errors and 1,932 significant pairs. These patterns were robust to omitting any single biological replicate: leaving out each replicate (one preparation day containing all six conditions) in turn yielded PSI estimates correlated with the full analysis at r = 0.95– 0.99, and all 1,462 significant deviations retained their direction.

Within-protein comparisons provided direct evidence that phosphoform organization could not be explained by parent-protein enrichment alone. Among 945 proteins with multiple quantified phosphosites, 528 (56%) exhibited PSI variation exceeding measurement error, including 64 (6.8%) with within-protein ranges greater than 2.0 log_2_ (**Supplementary Fig. 6b**). Because phosphosites on the same protein share a common parent-protein enrichment term, this heterogeneity cannot arise from a uniform shift in parent-protein proximity.

Talin-1 illustrates how spatial phosphoform preferences can differ from parent-protein representation (**Figs. 3b,f**; **Supplementary Fig. 6c**). Although abundant within CD18-labeled material, talin-1 was relatively under-represented compared with CAAX (−0.61). Its quantified phosphosites nevertheless spanned 1.83 log_2_ of PSI, with T144 at +2.00, S446 at +1.34 and S425 at +0.17, nearly coincident with the empirical center (+0.167). Lck showed similar within-protein heterogeneity, with five quantified phosphosites spanning 1.78 log_2_ from S121 (+0.23) to Y470 (+2.01). Kindlin-3 exhibited the complementary pattern: its parent protein was preferentially represented near CD18 (+0.88), whereas S484 showed lower representation than predicted by parent-protein enrichment (PSI −0.91). This relative under-representation persisted following ICAM-1 engagement (−0.58) and ICAM-1+PMA stimulation (−0.70; **Fig. 3g**). Immunoblotting detected talin-1 and kindlin-3 in plasma-membrane fractions from both cell lines at rest and after activation (**Fig. 3h**), supporting the distinction between gross membrane localization and receptor-proximal enrichment. Manual inspection of MS2 spectra supported the T144 and S484 phosphosite assignments.

Spatial phosphoform preferences also differed among residue and sequence classes. Before parent correction, phosphotyrosine sites showed greater CD18/CAAX enrichment (median +1.24; 90% above zero) than phosphoserine/threonine sites (median −0.12; 42% above zero; P = 1.1 × 10⁻²⁷ across 81 phosphotyrosine sites from 63 proteins). This difference persisted when each parent protein contributed one summary value (+1.22). Because phosphotyrosine-specific enrichment was not performed, differential recovery between residue classes limits quantitative comparisons. Nevertheless, after parent-protein correction, phosphotyrosine sites remained shifted above the empirical PSI center, whereas proline-directed Ser/Thr-Pro sites were distributed near it (**Supplementary Fig. 7c**). Thus, the relative phosphotyrosine preference was not explained solely by parent-protein enrichment, although these measurements do not establish absolute phosphorylation stoichiometry.

Together, these results demonstrate that the resting CD18 territory contains spatially heterogeneous phosphorylated forms of proteins whose representation cannot be predicted from parent-protein proximity alone.

### Sustained activation remodels phosphorylation while largely preserving protein and phosphoform organization

We next asked whether sustained activation reorganizes the protein and phosphoform organization established at rest. Comparing BSA with ICAM-1+PMA within CD18-labeled material, 60 of 6,456 proteins (0.93%) changed significantly (q < 0.05 and |log_2_ fold change| ≥ 0.5), compared with 1,434 of 4,627 phosphosites (31.0%), a 33-fold difference (**Figs. 4a**–**c**). Both measurements originated from the same labeling reactions and TMT multiplex, and the within-CD18 comparisons were independent of baseline differences between the CD18 and CAAX cell lines. The distinction was also evident without significance thresholds. Protein abundance was highly correlated between resting and activated conditions (Pearson r = 0.995; median absolute change 0.055 log_2_), whereas phosphosites showed greater variation (r = 0.879; median absolute change 0.330 log_2_; **Figs. 4d,e**). The functional-class organization and relative enrichment patterns of the resting CD18 territory were also broadly preserved after activation (**Supplementary Figs. 2b,c** and **3b,d**).

**Fig. 4.**
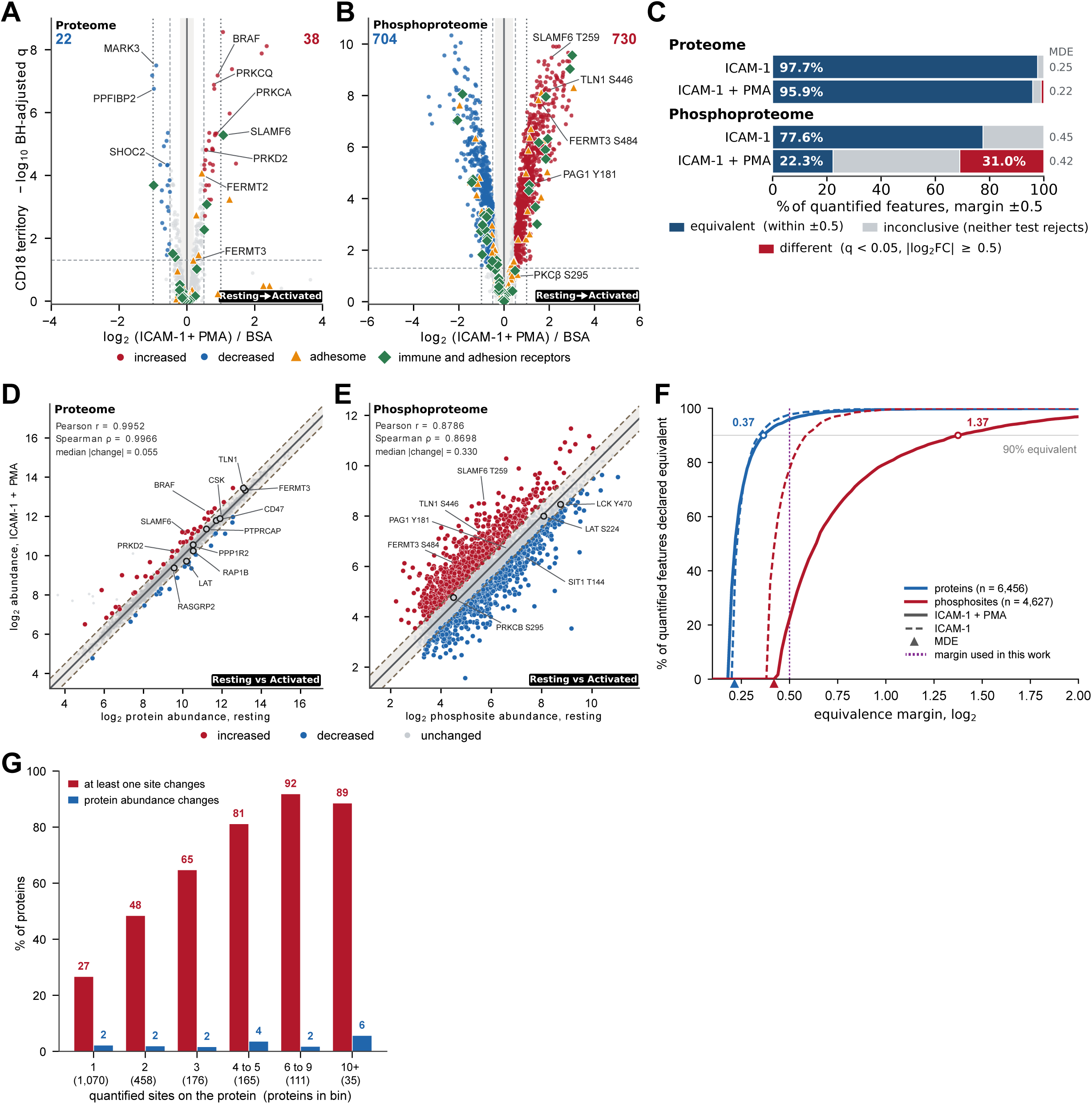
Sustained activation broadly remodels phosphorylation but largely retains protein architecture. **a,b,** Activation-associated changes in proteins (a) and phosphosites (b) in the CD18 sample, comparing resting BSA with ICAM-1+PMA at 30 min. Horizontal axes show log₂ fold change and vertical axes show −log₁₀ Benjamini–Hochberg-adjusted q. Red and blue denote increases and decreases meeting q < 0.05 and |log₂ fold change| ≥ 0.5; green diamonds and orange triangles mark immune and adhesion receptors and a curated adhesion-signaling protein set, respectively. The dashed horizontal line marks q = 0.05, dashed and dotted vertical lines mark ±0.5 and ±1 log₂, and the gray band marks the minimum detectable effect. In all, 38 proteins increased and 22 decreased, compared with 730 increased and 704 decreased phosphosites. **c,** Percentages of proteins and phosphosites formally equivalent within ±0.5 log₂, different by the statistical and effect-size thresholds, or inconclusive after ICAM-1 alone or ICAM-1+PMA. The minimum detectable effect (MDE) for each comparison appears in the right-hand column. **d,e,** Resting against activated abundance in proteins (d) and phosphosites (e); red, increased; blue, decreased; gray, unchanged by the stated criteria. Dashed lines mark ±0.5 log₂ relative to identity. Pearson r = 0.9952 and median absolute change 0.055 log₂ for proteins; Pearson r = 0.8786 and median absolute change 0.330 log₂ for phosphosites. **f,** Fraction of features meeting both one-sided equivalence tests at each symmetric margin; solid curves compare BSA with ICAM-1+PMA and dashed curves compare BSA with ICAM-1 alone. Triangles mark the MDEs given in c and the vertical dotted line marks the ±0.5 log₂ margin used for classification. Ninety percent of proteins were equivalent by approximately ±0.37 log₂, versus ±1.37 log₂ for phosphosites. **g,** Fraction of parent proteins with at least one activation-responsive phosphosite, or a protein-abundance response, by number of quantified sites per protein; bin labels give the number of parent proteins. Measurements derive from three biological replicates.

To determine whether the limited protein changes provided positive evidence for compositional stability, we performed two one-sided equivalence tests using a prespecified ±0.5 log_2_ margin, matching the differential-change effect-size threshold^28^. Of 6,456 proteins, 6,192 (95.9%) were formally equivalent between resting and activated CD18 samples within this interval, whereas 60 (0.9%) met the differential-change criterion and 204 remained inconclusive (**Fig. 4c**). The protein-level minimum detectable effect was 0.215 log_2_. Equivalence was robust across a range of margins, with 90% of proteins reaching equivalence at approximately ±0.37 log_2_ (**Fig. 4f**). Thus, compositional stability was supported by positive equivalence evidence between sustained endpoints rather than an absence of statistically significant differences. These measurements do not establish molecular immobility or exclude transient recruitment.

Phosphosites showed a markedly different response. Only 22.3% were equivalent within the same ±0.5 log_2_ interval, whereas 31.0% changed significantly and 46.7% remained inconclusive (**Fig. 4c**). Ninety percent equivalence required a margin of approximately ±1.37 log_2_, compared with ±0.37 log_2_ for proteins (**Fig. 4f**). The phosphosite minimum detectable effect was larger than that for proteins (0.420 versus 0.215 log_2_), reflecting lower measurement precision that increased the inconclusive fraction and made significant changes more difficult to establish. This difference persisted when evaluated at the parent-protein level. Among proteins with one quantified phosphosite, 27% contained an activation-responsive site, increasing to nearly 90% among proteins with six or more sites, whereas parent-protein abundance changes remained infrequent across coverage groups (**Fig. 4g**; **Supplementary Fig. 2d**).

The two stimulation conditions further distinguished ligand engagement from sustained co-stimulation. Immobilized ICAM-1 alone produced little remodeling at 30 min, with no proteins and one phosphosite meeting the differential-change criteria; 97.7% of proteins and 77.6% of phosphosites were equivalent within ±0.5 log_2_ (**Supplementary Figs. 3e,g**). Addition of PMA produced extensive phosphosite remodeling with comparatively limited protein changes (**Supplementary Figs. 3f,h**).

We then asked whether the extensive phosphorylation changes also altered the spatial phosphoform organization observed at rest. Despite widespread phosphosite remodeling, resting and activated PSI values remained strongly correlated (r = 0.85 across 4,564 site-parent pairs), with the standard deviation of activation-associated PSI changes (0.32 log_2_) substantially smaller than that of PSI itself (0.57 log_2_). The empirical PSI center shifted by only +0.045 log_2_. Most phosphorylation responses were shared between the CD18 and CAAX labeling environments, with site-minus-parent activation changes correlated at r = 0.88, leaving the spatial preferences of most phosphoforms largely unchanged. Following adjustment for the shift in the PSI distribution center, 24 phosphosites showed significant changes in PSI (13 increased and 11 decreased), compared with 35 before center adjustment (**Supplementary Fig. 6d**). An additional 466 sites exhibited |ΔPSI| > 0.5 log_2_, although many of these changes were statistically unresolved. PKCβ S295 provided a notable example, shifting from PSI +1.27 at rest to −0.54 following activation despite under-representation of its parent kinase near CD18. Distinguishing the contributions of the two labeling environments requires analysis of their differential activation responses.

Together, these findings establish that sustained activation extensively remodels phosphorylation while largely preserving both protein composition and the spatial phosphoform organization present at rest.

### Receptor-preferential activation responses are dominated by phosphosites

Activation-associated changes measured within the CD18 environment may include responses shared across the plasma membrane. Indeed, ICAM-1+PMA produced extensive phosphosite remodeling and selective protein redistribution in both proximity-labeled samples. To distinguish these shared responses from those preferentially associated with CD18, we subtracted the activation-associated change at the CAAX reference from the corresponding change within CD18 for each protein and phosphosite (**Fig. 5**; **Supplementary Fig. 8a**). This receptor-preferential difference, the statistical interaction between labeling environment and activation condition, compares activation responses between the two labeling environments; it does not measure absolute recruitment to or binding at the receptor. The corresponding analysis of immobilized ICAM-1 alone identified no significant protein-level differences (minimum q = 1.000).

**Fig. 5.**
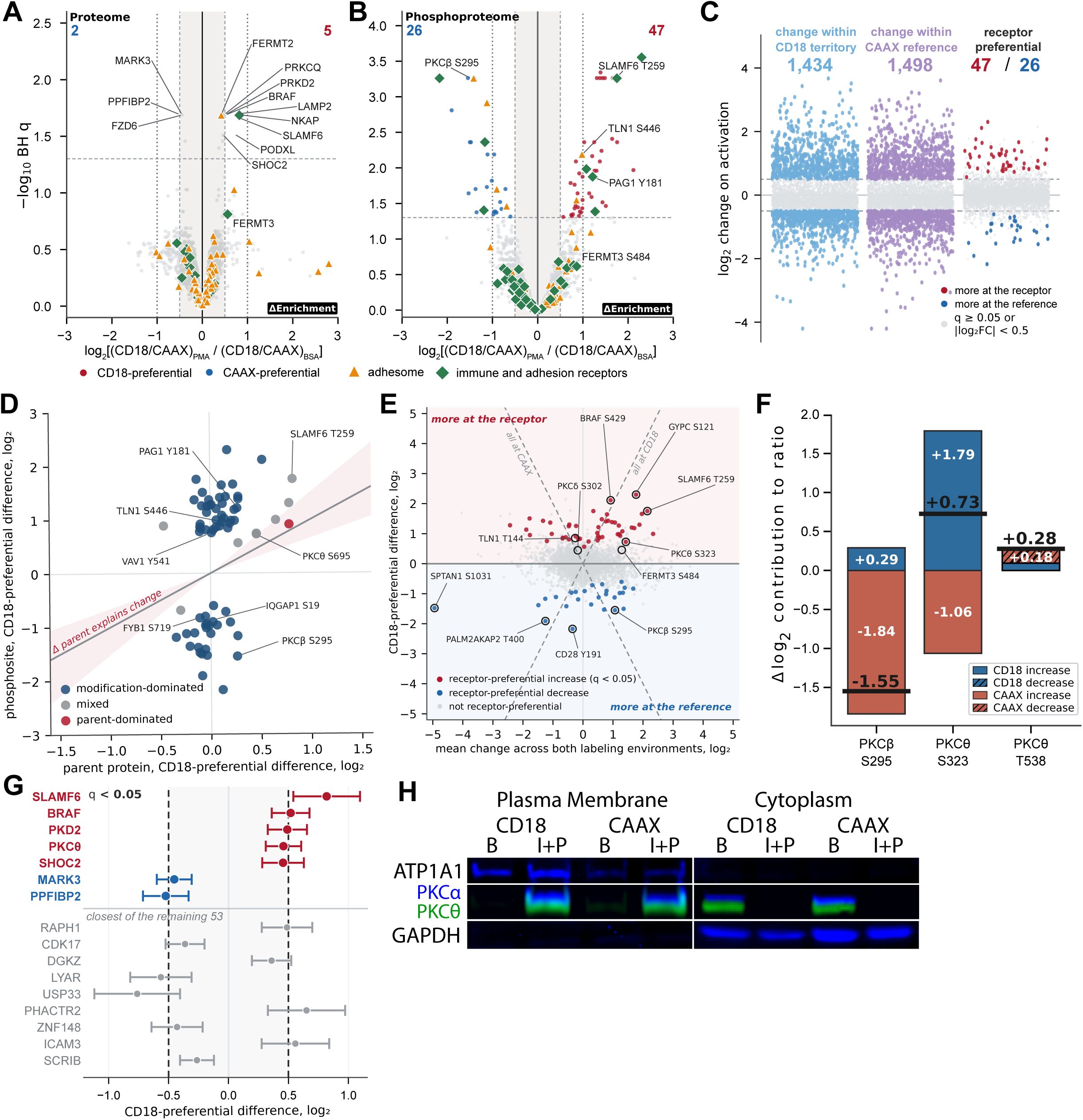
Receptor-preferential activation responses are dominated by phosphosites. **a,b,** Receptor-preferential protein (a) and phosphosite (b) activation differences, calculated as the change from BSA to ICAM-1+PMA in the CD18 samples minus the corresponding change at the CAAX reference. Horizontal axes show this difference, expressed as the change in log₂ CD18/CAAX enrichment, and vertical axes show −log₁₀ Benjamini–Hochberg-adjusted q. Red and blue denote relative responses toward CD18 and CAAX, respectively; these differences do not necessarily represent absolute recruitment. Green diamonds and orange triangles mark immune/adhesion receptors and a curated adhesion-signaling set. Colored points meet q < 0.05 and |difference| ≥ 0.5 log₂: five proteins had CD18-preferential and two CAAX-preferential differences, and the corresponding phosphosite counts were 47 and 26. Labels in a mark the seven proteins meeting both criteria, the additional proteins shown in g, and kindlin-2 and kindlin-3. **c,** Activation changes in phosphosites within CD18, within CAAX and in the receptor-preferential difference. The first two comparisons identified 1,434 and 1,498 changing sites, respectively; the receptor-preferential difference identified 47 sites relatively shifted toward CD18 and 26 toward CAAX. Gray denotes q ≥ 0.05 or |log₂ fold change| < 0.5. **d,** Receptor-preferential differences of the 72 significant phosphosites with quantified parent proteins plotted against the corresponding parent-protein differences. Classification as modification-dominated, mixed or parent-dominated follows the comparison of phosphosite and parent-protein differences; the diagonal denotes changes predicted from parent-protein redistribution alone. **e,** Receptor-preferential difference plotted against the mean activation change across both labeling environments; red and blue denote significant relative shifts toward CD18 or CAAX, and gray unresolved responses. Diagonal guides show changes confined to one labeling environment. **f,** Decomposition of PKC phosphosite differences into changes within CD18 and CAAX; the CAAX term enters with the opposite sign. Black bars show the net difference. **g,** Receptor-preferential differences and uncertainty for selected proteins with changes within CD18. Colored estimates met q < 0.05; dashed vertical lines indicate ±0.5 log₂. **h,** Immunoblot of PKCα and PKCθ in plasma-membrane and cytoplasmic fractions from CD18-APEX2 and CAAX-APEX2 cells under resting BSA (B) or ICAM-1+PMA (I+P) conditions, with ATP1A1 and GAPDH as plasma-membrane and cytoplasmic markers. PKCα and PKCθ were detected simultaneously in separate fluorescence channels. Both isoforms moved from the cytoplasm to the plasma-membrane fraction after stimulation in both cell lines, whereas proximity labeling resolved a CD18-preferential response for PKCθ but not PKCα. Representative of [n] independent experiments. Proteomic measurements derive from three biological replicates; q values are Benjamini–Hochberg adjusted.

The receptor-preferential protein response was substantially smaller than the remodeling measured within CD18. Among the 60 proteins that changed within CD18, the median absolute receptor-preferential difference was 0.25 log_2_, compared with a median absolute within-CD18 change of 0.67 log_2_ (**Fig. 5a**). Across all 6,456 quantified proteins, 16 receptor-preferential differences reached q < 0.05, of which seven also exceeded the prespecified absolute effect-size threshold of 0.5 log_2_. Only three of these seven proteins (BRAF, PPFIBP2 and SLAMF6) were among the 60 within-CD18 responders. Of the other four, FZD6 and PODXL changed within CD18 by less than the effect-size threshold, whereas LAMP2 and NKAP had differences driven mainly by changes at the CAAX reference. Accordingly, receptor-preferential differences were tested across the complete quantified proteome rather than restricted to proteins identified as changing within CD18.

Comparison with the spatial membrane reference also altered the interpretation of several canonical β2 integrin-associated proteins (**Supplementary Fig. 8b**). RASA3 increased by +0.74 log_2_ within CD18 (q < 0.05), but its receptor-preferential difference was smaller (+0.31) and statistically unresolved. RIAM also changed within CD18, yet its activation responses in the two labeling environments were formally equivalent within ±0.5 log_2_. Receptor-preferential differences for lamellipodin, ICAM-3, L-selectin and RGS10 remained inconclusive. These estimates were less precise than within-sample comparisons, partly because a down-weighted activated CAAX channel increased the median standard error from 0.071 to 0.098. Thus, the unresolved differences cannot establish either receptor preference or equivalence, whereas the RIAM result supports a shared activation response within the prespecified margin.

Phosphosites exhibited substantially more resolved receptor-preferential responses than proteins (**Figs. 5b,c**; **Supplementary Fig. 8a**). Of 4,627 quantified phosphosites, 1,434 changed within CD18 and 1,498 changed within CAAX. Seventy-three phosphosites met the receptor-preferential criteria (q < 0.05 and |difference| ≥ 0.5 log_2_), with 47 showing greater relative responses at CD18 and 26 at CAAX. Of these, 51 also changed significantly within CD18, whereas the remaining 22 were identified through their differential responses at the membrane reference. By comparison, only seven proteins met the same receptor-preferential criteria. The minimum detectable effect was larger for phosphosite differences than for protein differences (0.596 versus 0.295 log_2_), indicating that greater phosphosite measurement precision did not account for the larger number of resolved phosphosite responses.

Most receptor-preferential phosphosite responses were not explained by corresponding changes in parent-protein proximity. Of the 73 phosphosites, 72 had quantified parent proteins, 66 arose from proteins without a significant parent-protein difference, and 63 were classified as modification-dominated (**Fig. 5d**). Talin-1 S446 illustrates this distinction. Its activation-associated increase was +1.65 log_2_ within CD18 compared with +0.67 at CAAX, producing a receptor-preferential difference of +0.98 (q = 0.0065). In contrast, the parent protein (+0.21, q = 0.52) and S425 (+0.11, q = 0.90) showed no statistically resolved differences (**Supplementary Fig. 8c**). Other receptor-preferential phosphosite responses included PAG1 Y181 (+1.22), SLAMF6 T259 (+1.75), ADAP S719 (−0.83), GYPC S121 (+2.30) and CD28 Y191 (−2.17; **Fig. 5e**). These differences identify phosphoform-specific remodeling beyond that predicted by parent-protein redistribution, without establishing changes in absolute phosphorylation occupancy.

Protein kinase C (PKC) family members further illustrated the distinction between broad membrane responses and receptor-preferential remodeling. PKCθ S323 showed a positive receptor-preferential difference (+0.73 log_2_, q = 0.016), reflecting greater activation-associated gain at CD18, whereas PKCβ S295 showed a negative difference (−1.55, q = 5.5 × 10⁻⁴), reflecting greater gain at CAAX (**Fig. 5f**). PMA increased the proximity of all quantified phorbol-responsive PKC/PKD proteins within CD18, but only PKCθ and PKD2 showed significantly greater responses than at CAAX (**Fig. 5g**; **Supplementary Figs. 8d–f**). PKCα increased within CD18 but exhibited little differential response (+0.05, q = 0.904), whereas atypical PKCι changed little. Membrane fractionation showed that ICAM-1+PMA increased the plasma-membrane representation of both PKCα and PKCθ in CD18-APEX2 and CAAX-APEX2 cells (**Fig. 5h**). Despite this shared membrane redistribution, proximity labeling resolved a CD18-preferential increase for PKCθ but not PKCα, distinguishing general membrane recruitment from receptor-preferential organization.

Together, these results establish that the extensive phosphorylation remodeling measured near CD18 during sustained activation was largely shared with the spatial membrane reference. The smaller receptor-preferential response was dominated by phosphosites, most of which exhibited changes not explained by corresponding parent-protein redistribution. These findings distinguish broad membrane-associated signaling from phosphoform-specific remodeling within the CD18 territory at the sustained activation endpoint.

## DISCUSSION

These findings establish that the resting CD18 signaling environment is organized at two distinct levels: protein composition and the spatial representation of phosphorylated forms relative to their parent proteins. Receptors, docking adaptors and Rap GTPases were preferentially represented relative to several catalytic classes, while parent-protein correction revealed phosphoform differences not predicted by protein proximity. These differences included divergent phosphosites on the same protein (**Supplementary Fig. 6b**), which cannot be explained by uniform differences in parent-protein representation. Talin-1 illustrates this distinction. Although abundant in CD18-labeled material, talin was under-represented relative to CAAX, yet T144 and S446 showed strong parent-corrected preference near CD18 while S425 lay near the PSI distribution center. Conversely, kindlin-3 protein was enriched, whereas S484 was under-represented relative to its parent. Thus, protein proximity alone cannot resolve spatial phosphoform organization, and phosphosite enrichment without parent correction can mistake parent-protein representation for phosphoform preference. These observations indicate that receptor-proximal protein composition alone provides an incomplete description of the molecular states available for signaling.

Protein composition and spatial phosphoform organization were both established at rest and largely preserved at the sustained activation endpoint. At 30 min, 95.9% of proteins measured within the CD18 sample were equivalent within ±0.5 log_2_, whereas 31.0% of phosphosites changed in abundance. Nevertheless, resting and activated PSI remained correlated (r = 0.85), indicating that extensive phosphorylation remodeling did not produce a comparable reorganization of spatial phosphoform preferences. Most phosphorylation responses were shared between CD18 and CAAX, leaving a smaller receptor-preferential component dominated by phosphosites (73 versus seven proteins), with 63 of 72 phosphosites having quantified parents classified as modification-dominated. Among phorbol-responsive PKC/PKD proteins, only PKCθ and PKD2 showed greater activation-associated proximity increases at CD18 than at CAAX, whereas atypical PKCι changed little, consistent with its lack of a phorbol-responsive C1 domain^29,30^. These findings suggest that receptor-proximal signaling can be regulated through changes in the molecular states of locally available proteins, while much of their spatial organization is retained.

Similar organization may extend beyond β2 integrins. Pre-existing signaling environments have been described around CD3ζ, Lck and CD19, with CD19 also undergoing extensive phosphosite remodeling following activation^31–33^. However, receptor systems differ in the extent of activation-associated protein remodeling. GPER1 showed little compositional change, whereas CD19, angiotensin and adrenergic receptors exhibited changes associated with protein recruitment or trafficking^13,33–35^. In cardiomyocytes, β-adrenergic stimulation removed Rad from the CaV1.2 neighborhood through PKA phosphorylation, showing that a change in molecular state can itself reorganize local composition ^36^; stimulus-dependent redistribution has likewise been tracked around TRPV2 channels and within primary cilia ^37,38^. Differences in receptor type, stimulation and sampling time limit direct comparisons between these studies. Moreover, receptor-proximal enrichment may reflect general membrane residency, while phosphosite enrichment may follow the representation of its parent protein. Spatial membrane referencing, matched protein and phosphosite measurements, and parent correction distinguish these contributions, while receptor-minus-reference activation comparisons separate shared membrane responses from receptor-preferential remodeling^33,39^.

The organization observed at resting CD18 raises the possibility that rapid integrin responses are supported by signaling machinery already available near the receptor before stimulation. Leukocytes can arrest under vascular flow within fractions of a second^40–42^, requiring rapid responsiveness while limiting inappropriate adhesion. How much restraint is needed may depend on cell state: LFA-1 on effector T cells is spontaneously adhesive without inside-out signaling^43^, whereas circulating leukocytes must keep the receptor inactive until they encounter the right signal. In a receptor-proximal ready state, protein and phosphoform organization would precede stimulation, while biochemical and mechanical inputs regulate productive engagement. This framework provides a spatial starting point for established β2 integrin signaling through Rap1, talin, kindlin-3 and RIAM^5,7,44^. All quantified Rap isoforms were enriched near resting CD18, whereas CalDAG-GEFI was under-represented (−1.07), despite its capacity to associate with membranes^45,46^ and its requirement for phorbol ester-and chemokine-induced adhesion^6^. Functional importance therefore does not need to correspond to sustained receptor-proximal enrichment. Talin provides a complementary example. Phosphorylation at T144/T150 reduces talin membrane association and integrin activation, whereas preventing phosphorylation accelerates adhesion assembly^47,48^. The preferential representation of talin T144 relative to its parent protein could therefore reflect a locally available but functionally restrained pool, consistent with transient talin engagement at αLβ2^11^. Kindlin-3 S484 has also been implicated in thrombosis and hemostasis^49^, consistent with phosphorylation-dependent regulation of kindlin function^50^. Its increase after PMA was shared by both labeling environments, consistent with membrane-wide regulation of kindlin-3. Together, these examples are consistent with a pre-organized regulatory state near resting CD18, whose contribution to the kinetics of leukocyte arrest can now be tested directly.

Several mechanisms could generate and maintain the phosphoform organization observed at resting CD18. Docking adaptors were enriched relative to several of their reported regulatory enzymes, including Csk (−0.87) and selected phosphatases, while Ser/Thr phosphatase catalytic subunits were less enriched than activity-restraining regulators **(Supplementary Figs. 4a,c**). A PAG1 phosphopeptide spanning its Csk-docking region remained enriched and nearly invariant across conditions, although Csk itself was under-represented (**Supplementary Figs. 9a,b**)^24^. Independent single-molecule imaging also found no above-chance co-clustering of resting PAG and Csk nanoclusters^51^. Together, these observations suggest that locally available signaling substrates and phosphorylated docking platforms do not necessarily coincide with sustained enrichment of their associated regulatory enzymes. This asymmetry raises the possibility that receptor-proximal phosphoform organization can be maintained without continuous enzyme residence, through transient catalytic encounters, differences in enzyme access, phosphorylation turnover or regulated substrate accessibility. Selected kinase phosphoforms also exhibited spatial preferences relative to their parent proteins (**Supplementary Fig. 10**), and phosphotyrosine sites remained preferentially represented near CD18 after parent correction (**Supplementary Fig. 7c**). However, APEX2 measures time-integrated proximity rather than catalytic flux, and class-dependent reference offsets complicate comparisons among regulatory enzymes (**Supplementary Figs. 7d,e**). Related spatial regulatory mechanisms include kinetic segregation of transmembrane phosphatases at immune contacts^52,53^ and recruitment of phosphatase catalytic subunits to mature adhesions^54,55^. Membrane anchoring may also contribute by increasing local availability of prenylated Rap proteins, phosphoinositide-binding kindlin-3 and transmembrane adaptors, facilitating interactions such as Rap1B binding to talin F0^56^. Because enrichment was measured against a membrane-anchored reference, however, the receptor-proximal protein and phosphoform preferences observed here point to organization beyond membrane association alone.

Mechanical coupling may provide a complementary level of regulation that determines whether a pre-organized receptor environment supports force-bearing adhesion. The molecular clutch is the chain of cytoplasmic adaptor proteins that links ligand-engaged integrins to the moving actin cytoskeleton, so that intracellular forces are transmitted across the receptor to its ligand^57,58^. Actin-dependent coupling regulates integrin organization and activation: ligand-bound LFA-1 aligns with retrograde actin flow and tilts under intracellular loading^59^, ligand-engaged integrins in focal adhesions are co-aligned through a talin-mediated linkage to F-actin^59^, and myosin light chain phosphorylation supports integrin activation downstream of calcium entry^60^. Vinculin, Pyk2, paxillin, LIMS1 and ILK were detectable in CD18-labeled material at rest but showed little or negative enrichment relative to CAAX. Their presence without preferential enrichment is consistent with a distinction between local availability of signaling machinery and the formation of mature force-bearing adhesions, although proximity labeling does not establish whether these proteins are mechanically engaged. Biochemical readiness and mechanical engagement may therefore be regulated independently, allowing receptor-proximal signaling machinery to remain organized before formation of a productive, force-bearing adhesion.

Together, these findings show that the CD18 signaling environment is organized in both protein composition and phosphorylation state before activation, and that both are largely preserved at a sustained endpoint. The strategy that revealed this organization is not specific to integrins. A receptor-tethered proximity label, a compartment-matched membrane reference, and protein and phosphosite measurements from the same labeling reactions can be applied to any receptor that tolerates a cytoplasmic fusion. Parent correction and equivalence testing then turn a list of neighbors into four measurements: which proteins are preferentially represented near the receptor before stimulation, which of their phosphoforms are organized beyond what protein proximity predicts, whether stimulation rewrites local composition or molecular state, and which responses belong to the receptor rather than the surrounding membrane. Applied to antigen, chemokine and other rapidly acting receptors, the same design can determine whether pre-organized phosphoform states are a general feature of receptor signaling. If so, much of signaling may consist not of assembling machinery after stimulation but of changing the state of machinery already in place.

## ACKNOWLEDGEMENTS

We thank the Moore and Kalocsay Labs for helpful suggestions and insightful comments. Flow cytometry and cell sorting were performed at the UTHealth Houston Flow Cytometry Service Center. Fluorescence microscopy was performed at the Center for Advanced Microscopy, a Nikon Center of Excellence, in the Department of Integrative Biology & Pharmacology at McGovern Medical School, UTHealth Houston (RRID: SCR_025962).

## CONTRIBUTIONS

T.I.M. conceived the study. T.I.M. and N.H. designed the experiments. N.H. performed CRISPR gene editing and proximity labeling. H.J. performed the mass spectrometry experiments and mass spectrometry data analysis. T.I.M. and M.K. analyzed and interpreted the proteomics and phosphoproteomics datasets. T.I.M. wrote the original manuscript. N.H., H.J. and M.K. reviewed and edited the manuscript. All authors reviewed and approved the final version.

## METHODS

### Cell culture and cell lines

Jurkat T cells (clone E6-1) were obtained from the AmericanTissue Culture Collection (ATCC) and cultured in RPMI 1640 containing 2.05 mM L-glutamine and 11.10 mM glucose (Corning), supplemented with 2 mM L-glutamine (Corning), and 10% (v/v) heat-inactivated fetal bovine serum (FBS, Hyclone) and maintained at 37 °C in a humidified incubator with 5% CO₂

The CD18-APEX2 and CAAX-APEX2 cell lines used for proximity labeling were derived from the same parental Jurkat culture. For the CD18 bait line, endogenous *ITGB2* was disrupted by CRISPR-Cas9 and the knockout line was reconstituted with human CD18 carrying APEX2 at the C terminus of the cytoplasmic tail. The spatial membrane-reference line expressed CAAX-targeted APEX2.

### Generation of CD18-knockout Jurkat cells

The human *ITGB2* locus was disrupted using CRISPR-Cas9 with dual guide RNA’s [GTTCAACGTGACCTTCCGGC, TCGGCTGGCGCAACGTCACG] cloned into a modified version of px333 plasmid (a gift from Andrea Ventura, Addgene plasmid #64073) with a P2A-EGFP sequence appended to the C terminus of the Cas9 protein. Jurkat E6-1 cells were transfected by TransIT-Jurkat transfection reagent (Mirus Bio) using 2.5 µg of plasmid. 24 hours after transfection EGFP+ cells were isolated by fluorescence-activated cell sorting (FACS) (BD FACS Aria II SORP). Following recovery for 2 weeks, cells negative for surface CD18 were isolated by FACS using TS1/18 staining to the β-I domain of CD18. Following an additional recovery of two weeks, FACS was used again to enrich for CD18-negative cells.

Loss of CD18 protein was verified by immunoblotting (Cell Signaling Technology (CST), D4N5Z), and loss of surface CD18 (TS1/18) and CD11a (TS1/22) was confirmed by flow cytometry.

### Generation of stable CD18-APEX2 and CAAX-APEX2 cell lines

Human CD18 was fused at the C terminus of its cytoplasmic tail to APEX2 through a 14-residue linker (GGASGGSGSGPVAT). The spatial membrane-reference was constructed by fusing an 8 amino acid linker, PAGCMSCK, and CaaX sequence from HRas, CVLS, to the C terminus of APEX2. The Gateway system (ThermoFisher) was used to create lentiviral constructs of APEX2 fusions of CD18, and CAAX both followed by IRES-EGFP. Constructs were inserted into the pLX302 lentivirus plasmid (Addgene plasmid #25896 from David Root). 293T cells cultured in DMEM supplemented with 10% FBS were co-transfected with the lentiviral plasmid, psPAX2 and CMV-VSV-G using jetOPTIMUS DNA Transfection Reagent (Polyplus) to produce virus. Virus in supernatants was used to transduce Jurkat cells selected using 3 µg/mL puromycin (pLX302).

Stable populations were isolated by sequential fluorescence-activated cell sorting first by EGFP expression, then matching CD18 surface expression of CD18-APEX2 and CAAX-APEX2 with WT Jurkat cells. Surface expression of CD18 (TS 1/18) and CD11a (TS 1/22) was quantified by flow cytometry (BD LSRFortessa). Expression of CD18-APEX2 was verified by immunoblotting for CD18.

### Substrate preparation and cell stimulation

6-well plates were coated with 20 µg/mL Ni-NTA agarose purified soluble recombinant human ICAM-1-His6 (D1–D5)^61^ overnight at 4 °C. Surfaces were washed with DPBS (Corning) and blocked with 0.5% BSA in DPBS.

Cells were transferred to L15 media supplemented with 0.5% BSA and 3 mM biotinyl tyramide (Chemodex), applied at 4E6 cells/well, and incubated for 30 min at 37 °C under three conditions: BSA, immobilized ICAM-1, or immobilized ICAM-1 with 100 ng/mL phorbol 12-myristate 13-acetate (PMA, AdipoGen). BSA served as the unstimulated, non-adhesive baseline. 8E6 cells were used for each sample.

Three independent proximity-labeling experiments were performed, with each complete set of bait-by-condition samples prepared and processed on a separate day.

### Fractionation and whole cell lysate collection

Cell fractions were collected after treatment with PMA (100 ng/mL) and ICAM-1 (20 µg/mL) or BSA alone using Minute™ Plasma Membrane Protein Isolation and Cell Fractionation Kit (Invent, SM-005). 40E6 cells/sample were washed one time with DPBS and incubated in L15 media + 0.5% BSA with or without PMA. Cells were added to 6-well plates with or without ICAM-1 coating and spun down at 200 × *g* for 2 min prior to incubating for 30 min at 37 °C. After incubation, media was aspirated, cells were quickly resuspended in ice-cold DPBS and placed on ice. Cells were spun down at 4 °C and washed one time with ice-cold DPBS before proceeding to manufacturers fractionation protocol, performed in a 4 °C cold room. Final cytosolic fractions were suspended in a 1:1 ratio of DPBS and kit “Buffer A”. Plasma membrane fraction was resuspended in 25 µl resuspension buffer (8 M urea, 100 mM sodium phosphate, pH 8, 1% SDS, 100 mM ammonium bicarbonate).

Whole cell lysate was collected from cells treated as above (PMA+ICAM-1 or BSA on 6-well plates) using the Minute™ Total Protein Extraction Kit (Invent, SD-001/SN-002). 8E6 cells/sample were washed one time with DPBS and lysed in 100 µL of the kit’s Denaturing Buffer before running through kit column.

### Western blotting analysis

Biotin labeling was verified pre-and post-pulldown by western blot. 5% (pre-pulldown) and 10% (post-pulldown) of total sample was resolved on 7.5% Tris-glycine gels (Bio-Rad) and transferred to PVDF membrane (Bio-Rad) at 30 V for 16 hours in a 4 °C cold room. Membranes were blocked for 1 hour with 5% BSA in TBST (0.1% Tween-20) and incubated for 1 hour at RT with Streptavidin-HRP (CST, 3999) diluted at 1:5000. Membranes were washed 3 times with TBST and developed with Signal Fire™ ECL substrate (CST, 74709 & 46935) before being imaged (Azure Biosystems).

Cell fractions and whole cell lysate were analyzed by western blot for localization and expression of proteins of interest after treatment with ICAM-1 (20 µg/mL) + PMA (100 ng/mL) or BSA control. Protein concentration of samples was quantified by BCA assay. 10 µg from plasma membrane fractions and 20 µg from cytoplasmic fractions and whole cell lysate were resolved on 4-15% Tris-glycine gradient gels (Bio-Rad). Transfer and blocking was performed as described above. After blocking, membranes were incubated with appropriate primary antibodies at 1:1000 dilution for 1 hour at RT or overnight at 4 °C. The following primary antibodies were used: GAPDH (CST, 97166), ATP1A1 (Thermo Fisher, MA3-929), ITGB2 (CST, 73663), ITGAL (R&D Systems, MAB35952100), SHP-1 (CST, 26516), FERMT3 (CST, 10459), TLN1 (CST, 4021), PRKCQ (CST, 13643), FYN (Thermo Fisher, PA5-82851). Membranes were then washed 3 times and incubated with HRP-conjugated mouse (CST, 7076) and rabbit (CST, 7074) secondary antibodies at 1:5000 for 1 hour at RT. Membranes were washed 3 times before treatment with SignalFire HRP substrate and imaged.

### Fluorescent immunoblotting

For two-color detection of PKCα and PKCθ, cytoplasmic (20 µg) and plasma-membrane (10 µg) fractions were resolved as described above and transferred overnight at 25 V and 4 °C to low-fluorescence PVDF membrane (Azure Biosystems). Membranes were blocked for 10 min in Azure Fluorescent Blot Blocking Buffer (Azure Biosystems) and incubated with mouse anti-PKCα (Thermo Fisher, MA1-157) and rabbit anti-PKCθ (CST, 13643), together with mouse anti-ATP1A1 (Thermo Fisher, MA3-929) and mouse anti-GAPDH (CST, 97166) as plasma-membrane and cytoplasmic markers, each diluted 1:1,000 in blocking buffer. After three washes, membranes were incubated for 1 h at room temperature with goat anti-mouse AzureSpectra 550 (Azure Biosystems, AC2159) and goat anti-rabbit AzureSpectra 650 (Azure Biosystems, AC2165) secondary antibodies diluted 1:2,500 in blocking buffer. Membranes were then washed three times, dried and imaged on a Sapphire Biomolecular Imager (Azure Biosystems) in the 520-nm and 658-nm channels. PKCα, ATP1A1 and GAPDH were detected in the same channel and distinguished by molecular weight. Adjustments were applied linearly to each whole channel.

### APEX2 proximity labeling

Proximity labeling followed the peroxidase-catalyzed workflow of Kalocsay and colleagues^33,62–64^. Biotinyl tyramide (biotin-phenol) was added to the culture medium at a final concentration of 3 mM overnight before labeling. The following morning, cells were washed twice with DPBS and resuspended in L15 media supplemented with 0.5% BSA and 3 mM biotinyl tyramide, with or without PMA. After addition of cells to 6-well plates, cells were spun down for 2 min at 200 × g to adhere cells to the plate and placed at 37 °C. Following the 30-min stimulation period, proximity labeling was initiated by addition of H₂O₂ to a final concentration of 1 mM. A fresh H₂O₂ working solution was prepared immediately before each experiment by diluting a 30% (v/v) stock to 20 mM in PBS. Labeling proceeded for exactly 1 min.

The reaction was quenched by addition of ice-cold freshly prepared 2X quench buffer (20 mM sodium ascorbate, 10 mM Trolox, 20 mM sodium azide in PBS) directly to the medium. Cells were immediately spun down at 300 × *g* for 3 min, transferred into ice-cold quench buffer (10 mM sodium ascorbate, 10 mM sodium azide, and 5 mM Trolox in PBS), collected at 400 × *g* for 2 min, and washed again with cold quench buffer. Supernatant was removed, and cell pellets were flash-frozen in liquid nitrogen and stored at −80 °C until streptavidin enrichment.

Controls omitting biotinyl tyramide or H₂O₂ were processed in parallel and assessed by streptavidin blotting to verify APEX2-dependent biotinylation. All 18 TMTpro channels were assigned to the six bait-by-condition groups, so these controls were not carried as channels within the quantitative multiplex.

### Streptavidin enrichment of biotinylated proteins

All buffers were prepared fresh and filtered through 0.22-µm filters. Frozen cell pellets were lysed in 500 µl of lysis buffer (8 M urea, 100 mM sodium phosphate, pH 8.0, 1% (w/v) SDS, 100 mM ammonium bicarbonate, and 10 mM tris(2-carboxyethyl)phosphine (TCEP)). Lysates were incubated at room temperature for 10 min.

Proteins were precipitated by addition of an equal volume of ice-cold 55% trichloroacetic acid and incubated on ice for 15 min. Precipitates were collected at 17,000 × *g* for 10 min at 4 °C and washed four times with 500 µl of −20 °C acetone. Following the final wash, residual acetone was removed and pellets were air-dried on ice for 10 min.

Protein pellets were resuspended in 200 µl lysis buffer, sonicated for 10 min in a sonicator bath (Branson) and vortexed at room temperature for 1 hour. Insoluble material was removed by centrifugation at 20,000 × *g* for 10 min at room temperature. Proteins were alkylated with freshly prepared iodoacetamide, added from a 400 mM stock in 50 mM ammonium bicarbonate to a final concentration of 20 mM, for 25 min at room temperature in the dark. Alkylation was quenched by addition of dithiothreitol to a final concentration of 50 mM. Samples were then diluted with an equal volume of water to final concentrations of 4 M urea and 0.5% SDS.

For each sample, 75 µl of streptavidin magnetic-bead suspension (Pierce streptavidin magnetic beads, Thermo Fisher Scientific, 88817) was washed twice with 4 M urea, 0.5% SDS, and 100 mM sodium phosphate, pH 8.0, and added to the lysate. Samples were rotated overnight at 4 °C. Beads were subsequently washed three times with 4 M urea, 0.5% SDS, and 100 mM sodium phosphate, pH 8.0; then three more times with the same buffer lacking SDS. Beads were transferred to fresh tubes after the first wash of each series and stored at −80 °C until digestion.

### On-bead digestion and TMTpro labeling

Streptavidin-bound proteins were digested on-bead in 200 mM EPPS, pH 8.5, containing 2% acetonitrile. Samples were first digested for 3 h with endoproteinase LysC (Wako) at an enzyme-to-substrate ratio of 1:50 and then overnight with additional trypsin (Promega) at an enzyme-to-substrate ratio of 1:100, at 37 °C with gentle agitation. Digestion efficiency was assessed by mass spectrometry on a small aliquot of each reaction before TMTpro labeling.

Peptides from the 18 experimental samples were labeled with TMTpro 18-plex reagents (Thermo Fisher Scientific, A52045). Labeling was performed in 200 mM EPPS, pH 8.5, after adjustment to 30% (v/v) acetonitrile. Labeling efficiency was evaluated by mass spectrometry on a StageTip-purified aliquot prepared by pooling approximately 1% of each labeling reaction. TMTpro labeling efficiency exceeded 95%, as determined by searching TMTpro (+304.207 Da) as a variable modification on peptide N termini and lysine residues. Labeling reactions were quenched by addition of 0.5% hydroxylamine for 15 min and acidified with formic acid. Acetonitrile was removed by vacuum centrifugation.

Labeled peptides were resuspended in 1% formic acid and 0.1% trifluoroacetic acid (TFA) and desalted by gravity-flow reversed-phase chromatography using C18 Sep-Pak cartridges (Waters, WAT054945). Columns were conditioned sequentially with methanol, acetonitrile, and 1% formic acid containing 0.1% TFA. Samples were washed with three column volumes of 1% formic acid and eluted with 95% acetonitrile containing 1% formic acid. Eluted peptides were dried to near completion before phosphopeptide enrichment.

### Microscale phosphopeptide enrichment

TMTpro-labeled peptides were subjected to microscale Fe-NTA enrichment so that phosphopeptide and protein-level measurements originated from the same APEX2 labeling reactions and pooled peptide preparation.

Dried peptides were resuspended in 50 µl Fe-NTA binding buffer containing 0.1% TFA and 80% acetonitrile using the High-Select Fe-NTA Phosphopeptide Enrichment Kit (Thermo Fisher Scientific, A32992). Microscale enrichment columns were prepared by inserting a single C18 disk punched from an Empore C18 solid-phase extraction disk (Sigma-Aldrich, 66883-U) with a 16-gauge blunt-ended needle into a pipette tip. Tips were conditioned with 100 µl acetonitrile before addition of approximately 20 µl of Fe-NTA bead slurry, corresponding to approximately 10 µl of beads, transferred from a High-Select column. Buffers were passed through the tip by gentle air pressure so that the beads did not run dry.

Fe-NTA beads were washed twice with 70 µl binding buffer. Peptide samples were loaded onto the beads and incubated for 30 min, with the bead suspension mixed every 10 min and any air bubbles between the peptide solution and the beads removed by gentle pipetting. The unbound fraction was collected together with three 70-µl binding-buffer washes, one 70-µl ultrapure-water wash, and one 50-µl acetonitrile wash. The combined flow-through and washes constituted the non-phosphopeptide fraction used for protein-level proximity analysis and were dried by vacuum centrifugation before high-pH reversed-phase fractionation.

Bound phosphopeptides were eluted twice with 25 µl alkaline Fe-NTA elution buffer consisting of aqueous ammonia, pH 11.3, followed by 50 µl acetonitrile. Eluates were pooled directly in autosampler-vial inserts, dried completely by vacuum centrifugation, and resuspended in 6 µl of 1% formic acid for LC-MS/MS analysis.

### High-pH reversed-phase peptide fractionation

The combined non-phosphopeptide flow-through was fractionated by alkaline reversed-phase chromatography using the Pierce High-pH Reversed-Phase Peptide Fractionation Kit (Thermo Fisher Scientific, 84868) using a modified elution scheme of 12 sequential fractions (10, 12.5, 15, 17.5, 20, 25, 30, 35, 40, 50, 65, and 80% acetonitrile). The 12 fractions were dried by vacuum centrifugation, desalted by StageTip, and resuspended in 1% formic acid before LC-MS/MS analysis.

### Liquid chromatography–tandem mass spectrometry

Peptides were analyzed on an Orbitrap Ascend Tribrid mass spectrometer (Thermo Fisher Scientific) using FAIMS and MS2-based TMTpro quantification with TurboTMT enabled. Each of the 12 high-pH reversed-phase fractions was analyzed using a 120-min LC-MS/MS method with FAIMS compensation voltages of −40, −60, and −80 V. MS1 spectra were acquired in the Orbitrap at a resolution of 120,000 over a scan range of 350–1,400 m/z with a maximum injection time of 100 ms. Precursors with charge states of 2–6 were selected using a 2-s cycle time and a dynamic exclusion of 60 s. Precursors were isolated with a 0.7 m/z quadrupole isolation window and fragmented by HCD at a normalized collision energy of 33%. MS2 spectra were acquired in the Orbitrap at a resolution of 30,000 with TurboTMT enabled, a first mass of 120 m/z, and a maximum injection time of 250 ms.

The microscale Fe-NTA-enriched phosphopeptide sample was analyzed as a single 180-min LC-MS/MS run on the Orbitrap Ascend coupled to a Vanquish Neo nano-UHPLC system. Peptides were separated at 500 nL/min using a 25-cm, 75-µm-inner-diameter analytical column. The LC gradient reached 10% mobile phase B at 10 min, 27.5% B at 151 min, 42% B at 169 min, and 100% B at 176 min, followed by column washing through 180 min. FAIMS compensation voltages of −40, −60, and −80 V were used with a carrier gas flow of 4.5 L/min. MS1 spectra were acquired in the Orbitrap at a resolution of 120,000 over 375–1,500 m/z with a maximum injection time of 100 ms. Precursors with charge states of 2–8 were selected using a 3-s cycle time. Precursors were isolated with a 0.7 m/z quadrupole isolation window and fragmented by HCD at a normalized collision energy of 33%. MS2 spectra were acquired in the Orbitrap at a resolution of 30,000 with TurboTMT enabled, a first mass of 110 m/z, and a maximum injection time of 600 ms.

### Mass spectrometry database searching and quantification

Raw spectra were converted to mzXML with correction of monoisotopic peak assignments. Spectra were searched using the Comet search engine against a size-sorted forward and reverse human UniProt reference proteome database containing common contaminants (Human_Uniprotref_sorted_REV_contam_04192024.fasta). Searches specified trypsin, a precursor mass tolerance of 20 ppm, and a fragment-ion tolerance of 0.02 Da. Carbamidomethylation of cysteine (+57.0215 Da) and TMTpro modification of lysine residues and peptide N termini (+304.2071 Da) were specified as static modifications, and oxidation of methionine (+15.9949 Da) was specified as a variable modification. For phosphopeptide searches, phosphorylation of serine, threonine, and tyrosine (+79.9663 Da) was included as a variable modification. Peptide-spectrum matches were filtered by linear discriminant analysis against the target-decoy database to a false discovery rate of 1% at both the peptide-spectrum-match and protein levels, and peptides used for quantification were required to have an isolation specificity ≥0.70 and a summed reporter-ion signal-to-noise ≥200 across all channels.

### Protein and phosphosite filtering

The 18 TMTpro channels represented two labeling environments, CD18-APEX2 and CAAX-APEX2, under three conditions, BSA, ICAM-1, and ICAM-1 plus PMA, across three biological replicates on independent preparation days. Proteins were retained when quantified in all 18 channels and represented by at least two peptides. Of 7,286 protein groups, 7,281 had complete channel quantification and 6,456 remained after application of the two-peptide criterion. The median retained protein was represented by 10 peptides. Phosphosites were retained if quantified in all 18 channels, the same complete-channel rule applied to proteins, giving 4,627 of 8,832 site entries from 2,078 genes. Of the excluded entries, 4,173 were identified but quantified in no channel, because none of their peptide-spectrum matches passed the quantification filters (isolation specificity ≥ 0.70 and summed reporter signal-to-noise ≥ 200 across the 18 channels), and 32 were quantified in 15 to 17 channels. Because these filters act on the pooled spectrum, the first loss is common to all channels and cannot depend on bait or condition; consistent with this, among sites with a quantified parent protein, unquantified and retained sites had the same parent resting CD18/CAAX distribution (median −0.36 log2 for both; Mann-Whitney P = 0.69). Each retained entry is a quantified phosphopeptide form: 4,123 carry a single phosphorylation site and 504 are multiply phosphorylated forms treated as separate features. Most entries (3,509, 75.8%) were quantified from a single peptide-spectrum match, so no peptide-count requirement was applied at the site level.

## Data analysis

### Loading normalization

For each protein and phosphosite, per-channel sn_sum values were median loading-normalized. The median signal across positively quantified features was calculated separately for each of the 18 channels, and each channel was multiplied by a single factor that brought its median to the mean of the 18 channel medians.

### Linear modeling and channel-quality weights

Normalized protein and phosphosite abundances were modeled independently using a cell-means design containing six bait-by-condition groups: CD18 and CAAX under BSA, ICAM-1, and ICAM-1 plus PMA conditions. Preparation day was included as a blocking factor.

Per-channel quality weights were estimated iteratively from model residuals using an arrayWeights-style procedure. Weights were updated over six iterations using a damping exponent of 0.5, bounded between 0.1 and 5.0 during estimation, and normalized to a geometric mean of one. Two under-loaded day-1 CAAX channels, ICAM-1 and ICAM-1 plus PMA, received the lowest weights and were retained rather than excluded. Weighted least squares was used to estimate the six group means and preparation-day effects.

### CD18 enrichment and activation contrasts

CD18 proximity enrichment was defined within each condition as the fitted CD18 mean minus the corresponding CAAX mean on the log₂ scale: **log₂(CD18/CAAX).**

Positive values indicate preferential representation in the CD18 proximity environment relative to the spatial membrane reference; negative values indicate greater representation in the reference environment.

Activation effects were evaluated first as within-bait contrasts comparing ICAM-1 or ICAM-1 plus PMA with BSA separately in the CD18 and CAAX arms. Receptor-preferential remodeling was then quantified as the difference of differences: the activation-associated change in the CD18 arm minus the corresponding change in the CAAX arm.

### Phosphospatial index

Phosphosites were matched to quantified parent proteins by gene symbol, retaining only sites whose parent protein passed the protein-level filters. For each matched site, the phosphospatial index (PSI) was defined as: **PSI = log₂(site CD18/CAAX) − log₂(parent protein CD18/CAAX).**

A total of 4,564 site-parent pairs representing 2,015 genes were available for analysis.

Because the protein and phosphosite matrices were normalized independently, layer-wide offsets were not assumed to cancel. PSI was therefore interpreted relative to the empirical center of its distribution within each condition rather than relative to zero. The resting PSI median was +0.167 log₂, with alternative center estimates ranging from +0.135 to +0.217.

### Statistical analysis

Feature-level contrasts were tested using moderated *t* statistics after empirical-Bayes shrinkage of residual variances with an intensity-dependent mean-variance trend. The cell-means model contained 10 residual degrees of freedom before variance moderation. *P* values were two-sided unless otherwise stated and were adjusted for multiple comparisons within each contrast family using the Benjamini-Hochberg procedure.

Unless otherwise indicated, a protein or phosphosite was classified as changed when it satisfied both *q* < 0.05 and an absolute log₂ fold change of at least 0.5.

### Equivalence testing

Compositional stability was tested using two one-sided tests against a prespecified equivalence margin of ±0.5 log₂. Benjamini-Hochberg correction was applied across all quantified features. Features were classified as equivalent when the equivalence test reached *q* < 0.05, different when they met the differential-change criteria above, and inconclusive otherwise.

The equivalence margin was evaluated against the experiment’s null behavior. Simulation from fitted protein-level standard errors placed the 99th percentile of apparent null change at 0.205 log₂, while the empirical BSA-versus-ICAM-1 comparison placed it at 0.237 log₂. Equivalence results were additionally evaluated continuously across margins from 0.1 to 2.0 log₂.

Minimum detectable effects were calculated as the effect magnitude detectable with 80% power at a two-sided α of 0.05 using the median fitted standard error and moderated degrees of freedom of the corresponding contrast.

### Receptor-preferential difference testing

The receptor-preferential difference was evaluated as the statistical interaction between labeling environment and activation condition, that is, the change from BSA to ICAM-1+PMA within CD18 minus the corresponding change at the CAAX reference. Multiple-testing correction was applied across the complete quantified protein or phosphosite set rather than only to features first selected as changing within one arm.

### PSI statistical testing

For each site-parent pair, deviation of PSI from the empirical distribution center was tested using the site-minus-parent estimate and the propagated standard error of the two measurements. The variance calculation used the sum of the squared standard errors of the site and parent estimates. Benjamini-Hochberg correction was applied across the 4,564 site-parent tests. As a sensitivity analysis that does not assume independent site and parent errors, the same linear model was fitted directly to per-channel site-minus-parent log₂ differences for each matched pair, with re-estimated channel weights and empirical-Bayes moderation. PSI estimates agreed closely with the propagated values (Pearson r = 0.998), standard errors were smaller (median 0.144 versus 0.169 log₂), and 1,932 rather than 1,462 resting pairs differed from the distribution center at q < 0.05, indicating that the independent-error calculation is conservative.

### Leave-one-day-out sensitivity analysis

Robustness to individual preparation days was assessed by removing each day in turn and refitting the complete analysis to the remaining 12 channels, including re-estimation of channel-quality weights, empirical-Bayes moderation, multiple-testing correction, and PSI centering. Effect-size agreement rather than significance counts was used to evaluate stability because the reduced fits contain fewer residual degrees of freedom. Across the three reduced fits, resting CD18/CAAX enrichment correlated with the full fit at r = 0.980–0.994 for proteins and r = 0.974–0.992 for phosphosites (median absolute differences 0.05–0.07 and 0.07–0.11 log₂). Resting PSI correlated at r = 0.953–0.986 (median absolute difference 0.07–0.10 log₂), deviations of each site from the mean PSI of its protein at r = 0.951–0.983, and the empirical PSI center moved by at most 0.033 log₂. All 1,462 sites that deviated significantly from the center in the full fit kept the direction of deviation in every reduced fit. Activation-associated changes correlated at r = 0.825–0.921 for proteins and r = 0.983–0.989 for phosphosites, every phosphosite classified as changed kept its direction of change, and resting and activated PSI correlated at r = 0.76–0.86 within each reduced fit (0.85 in the full fit). Receptor-preferential differences were the least stable (r = 0.590–0.677 for proteins and 0.733–0.914 for phosphosites), although all seven proteins with a significant receptor-preferential difference in the full fit retained its sign in every reduced fit.

### Complete-channel sensitivity analysis

To test whether the complete-channel requirement shaped the phosphosite results, the 32 partially quantified sites were added back and each was fitted on its observed channels, with site-specific residual degrees of freedom carried into empirical Bayes moderation, as limma does for missing values. Normalization factors and array weights were those of the primary fit; the variance trend, prior degrees of freedom, Benjamini-Hochberg correction and PSI centers were re-estimated. No added site lacked all three replicates of any condition. The 42 missing values were split between the arms (23 CD18, 19 CAAX) and behaved as reporter dropout at low signal rather than absence from one bait: for 24 of the 32 sites every missing value fell in one of the two conditions with the lowest fitted signal for that site, and the median residual variance of these sites was 5.1-fold the intensity trend, against 1.0-fold for complete sites. The added sites were enriched for large activation changes (26 of 32 changed; median absolute change 2.46 against 0.33 log2 for retained sites). Including them lowered the phosphosite prior degrees of freedom from 8.09 to 7.46 and changed the headline results by less than half a percentage point: 31.3% of phosphosites (1,460 of 4,659) changed in the CD18 arm on activation, against 31.0% (1,434 of 4,627); 76 phosphosites were receptor-preferential, against 73; and 1,468 of 4,593 site-parent pairs (32.0%) deviated from the PSI center at rest, against 1,462 of 4,564 (32.0%), with the center unchanged at +0.167 log2. Resting and activated PSI correlated at r = 0.850, against 0.849. On the 4,627 common sites, estimates were identical and every activation change and receptor-preferential call was retained; eight PSI calls with primary q between 0.047 and 0.049 moved just above 0.05. The same rule added one protein group (7,281 of 7,286 groups were complete) and left the 60 activation-changed and seven receptor-preferential proteins unchanged. Because the excluded sites were enriched for large changes, the complete-channel requirement makes the phosphosite change fraction, and its 33-fold separation from the protein change fraction, slightly conservative rather than inflated.

## Data availability

The mass spectrometry proteomics data have been deposited to the ProteomeXchange Consortium via the PRIDE partner repository^65^ with the dataset identifier PXD085432. The deposition includes the raw files, search results and the quantified protein, peptide, phosphoprotein and phosphopeptide tables analyzed in this study.

