## Supplementary Material for "Receptor-proximal phosphoform organization precedes β2 integrin activation"

**This file includes: Supplementary Figures 1–10 and their legends.**

### Supplementary Figure 1

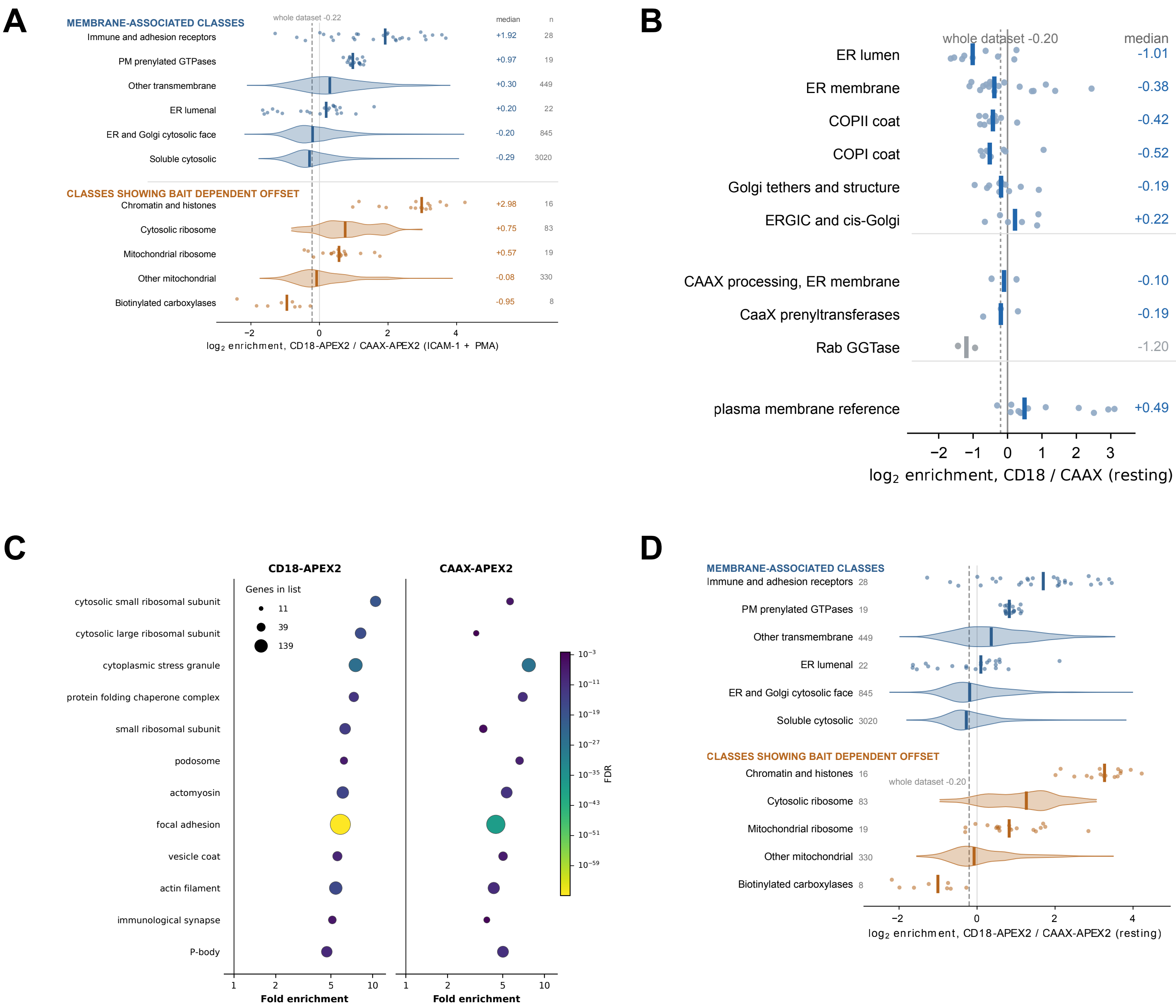

**Supplementary Fig. 1 | Spatial reference behavior of the CD18-APEX2 and CAAX-APEX2 proximity proteomes**

**a,** $\log_2$ CD18-APEX2/CAAX-APEX2 enrichment after ICAM-1+PMA activation across membrane-associated protein classes and classes revealing systematic differences between the two cell preparations. Each point represents one protein; violin plots show the class distributions, vertical bars denote class medians and the dashed line denotes the whole-dataset median (−0.22 $\log_2$). Class medians and protein numbers are given at right. **b,** Resting CD18/CAAX enrichment of proteins assigned to early secretory compartments, to CAAX-processing machinery and to a plasma-membrane reference set. Points represent individual proteins and vertical bars denote class medians; the dashed line denotes the whole-dataset median (−0.20 $\log_2$). Class medians and protein numbers are given at right. **c,** Gene Ontology cellular-component enrichment among proteins recovered with CD18-APEX2 (left) and CAAX-APEX2 (right). Symbol size indicates the number of contributing genes, color denotes false discovery rate and the horizontal axis shows fold enrichment over the proteome background. Both labeling environments recover the same categories, with focal adhesion the largest and most significant term in each. **d,** As in a for the resting condition (whole-dataset median −0.20 $\log_2$). CD18/CAAX estimates derive from three biological replicates; n values denote proteins, not replicates.

### Supplementary Figure 2

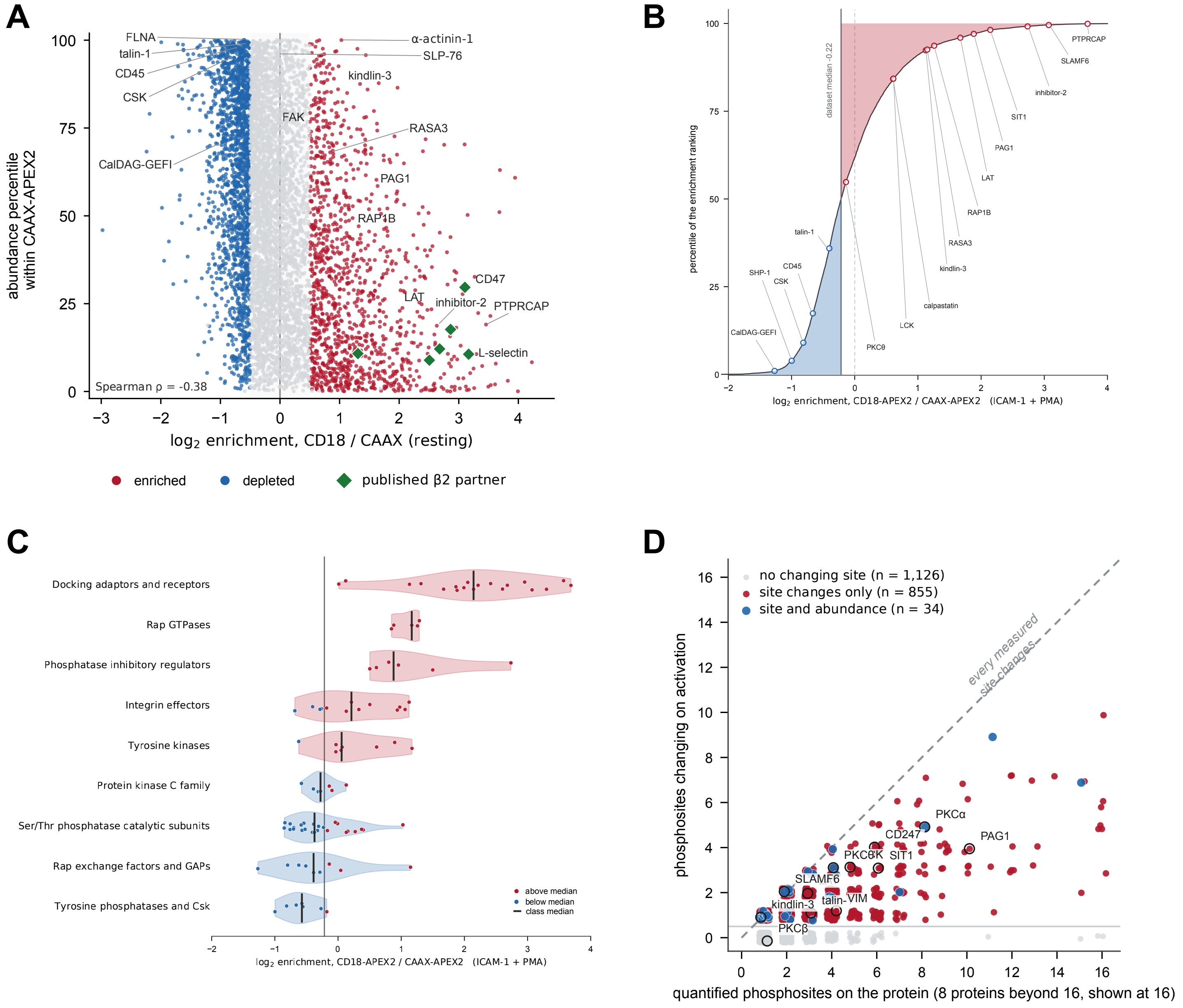

**Supplementary Fig. 2 | Abundance, functional class and phosphosite coverage of the CD18 proximity proteome**

**a,** Resting CD18/CAAX enrichment plotted against protein-abundance percentile within the CAAX-APEX2 reference. Red and blue denote proteins enriched or under-represented at CD18 according to the prespecified statistical and effect-size criteria; green diamonds mark previously reported β2 integrin-associated surface proteins. Ranking by abundance in the reference arm produces an apparent negative relationship (Spearman $\rho = -0.38$) that is expected from the arithmetic coupling of a ratio to the abundance term in its denominator (compare Fig. 2b). **b,** Position of selected signaling proteins across the activated (ICAM-1+PMA) CD18/CAAX enrichment ranking. The curve is the cumulative distribution of enrichment; red and blue shading mark proteins above and below the dataset median ($-0.22$ $\log_2$, dashed line). **c,** Activated (ICAM-1+PMA) CD18/CAAX enrichment distributions for prespecified functional classes. Points represent individual proteins and black marks denote class medians; red and blue indicate proteins above or below the dataset median. **d,** Number of quantified phosphosites per protein plotted against the number changing between resting and ICAM-1+PMA conditions within the CD18 samples ($q < 0.05$ and $|\log_2$ fold change$| \geq 0.5$). Proteins are classified as carrying no changing site (gray, n = 1,126), phosphosite changes only (red, n = 855) or both phosphosite and protein-abundance changes (blue, n = 34). The dashed diagonal is the expectation if every quantified site changed; proteins with more than 16 quantified sites are drawn at 16. Measurements derive from three biological replicates.

### Supplementary Figure 3

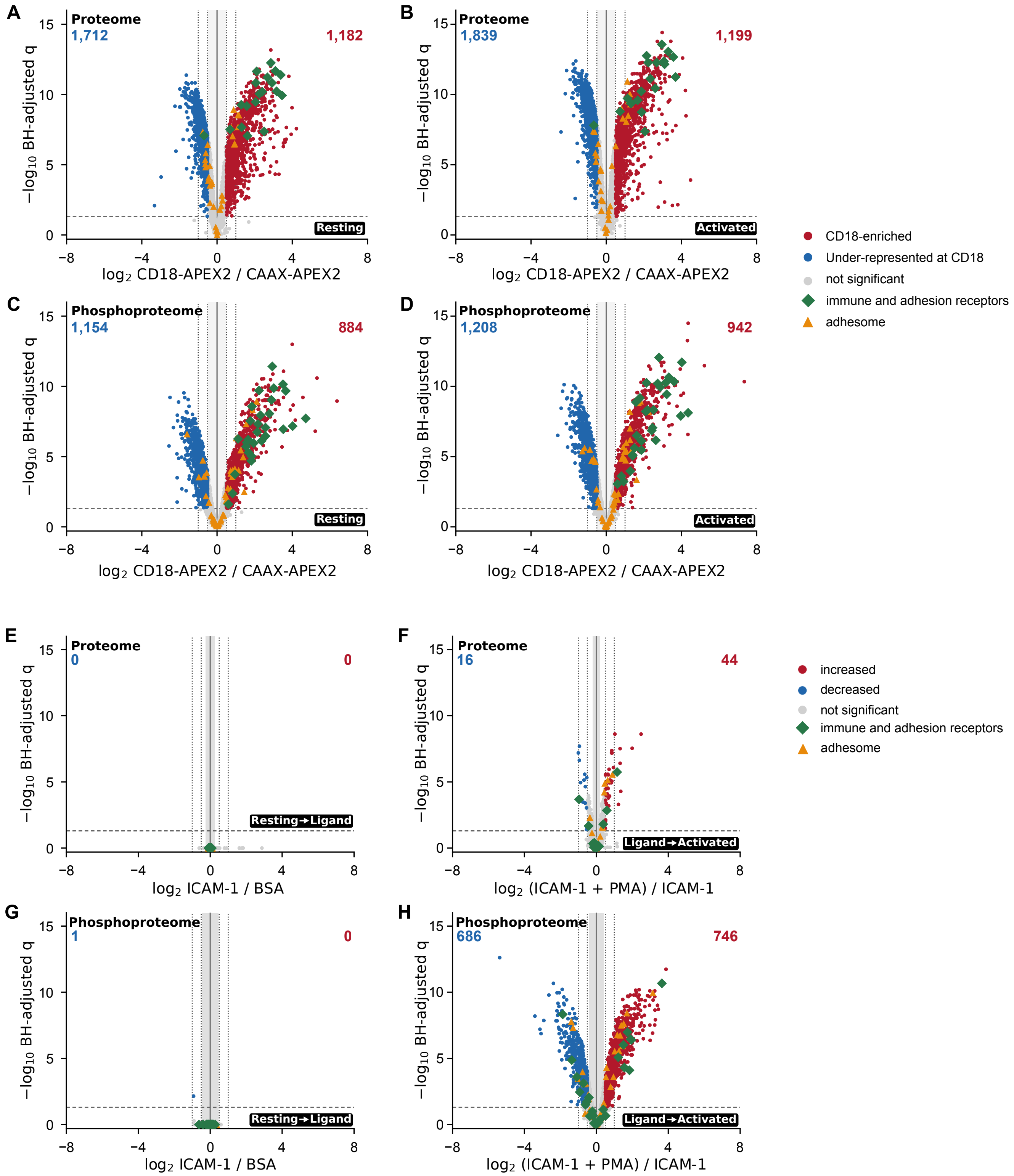

**Supplementary Fig. 3 | CD18/CAAX enrichment at the two endpoint states and the two sequential activation steps within the CD18 territory**

**a–d**, CD18-APEX2/CAAX-APEX2 enrichment of quantified proteins (a,b) and phosphosites (c,d) at rest (a,c) and after ICAM-1+PMA (b,d). Horizontal axes show $\log_2$ enrichment and vertical axes show $-\log_{10}$ Benjamini–Hochberg-adjusted q. Red and blue denote features enriched or under-represented at CD18 using $q < 0.05$ and $|\log_2$ fold change$| \geq 0.5$; counts are given in the corners. Of 6,456 proteins, 1,712 were under-represented and 1,182 enriched at rest, and 1,839 and 1,199 after activation; of 4,627 phosphosites, 1,154 and 884 at rest, and 1,208 and 942 after activation. Enrichment is a between-arm comparison and is subject to the class-dependent offsets shown in **Fig. 1e** and **Supplementary Fig. 1. e–h**, Activation-step changes within the CD18 samples for proteins (e,f) and phosphosites (g,h): resting BSA to ICAM-1 alone (e,g) and ICAM-1 alone to ICAM-1+PMA (f,h). Red and blue denote significant increases and decreases under the same criteria. Ligand presentation alone changed no protein (minimum q = 1.000) and one phosphosite (TRPM7 T1471); the subsequent PMA step decreased 16 and increased 44 proteins and decreased 686 and increased 746 phosphosites. In all panels the dashed horizontal line marks q = 0.05, dotted vertical lines mark ±0.5 and ±1 $\log_2$, and the gray band marks the minimum detectable effect at 80% power (a, 0.23; b, 0.20; c, 0.45; d, 0.40; e, 0.25; f, 0.21; g, 0.45; h, 0.41 $\log_2$). Green diamonds and orange triangles identify immune and adhesion receptors and the curated adhesion-signaling set, respectively. Moderated t-tests with Benjamini–Hochberg correction within each contrast; n = 6,456 proteins and 4,627 phosphosites from three biological replicates.

### Supplementary Figure 4

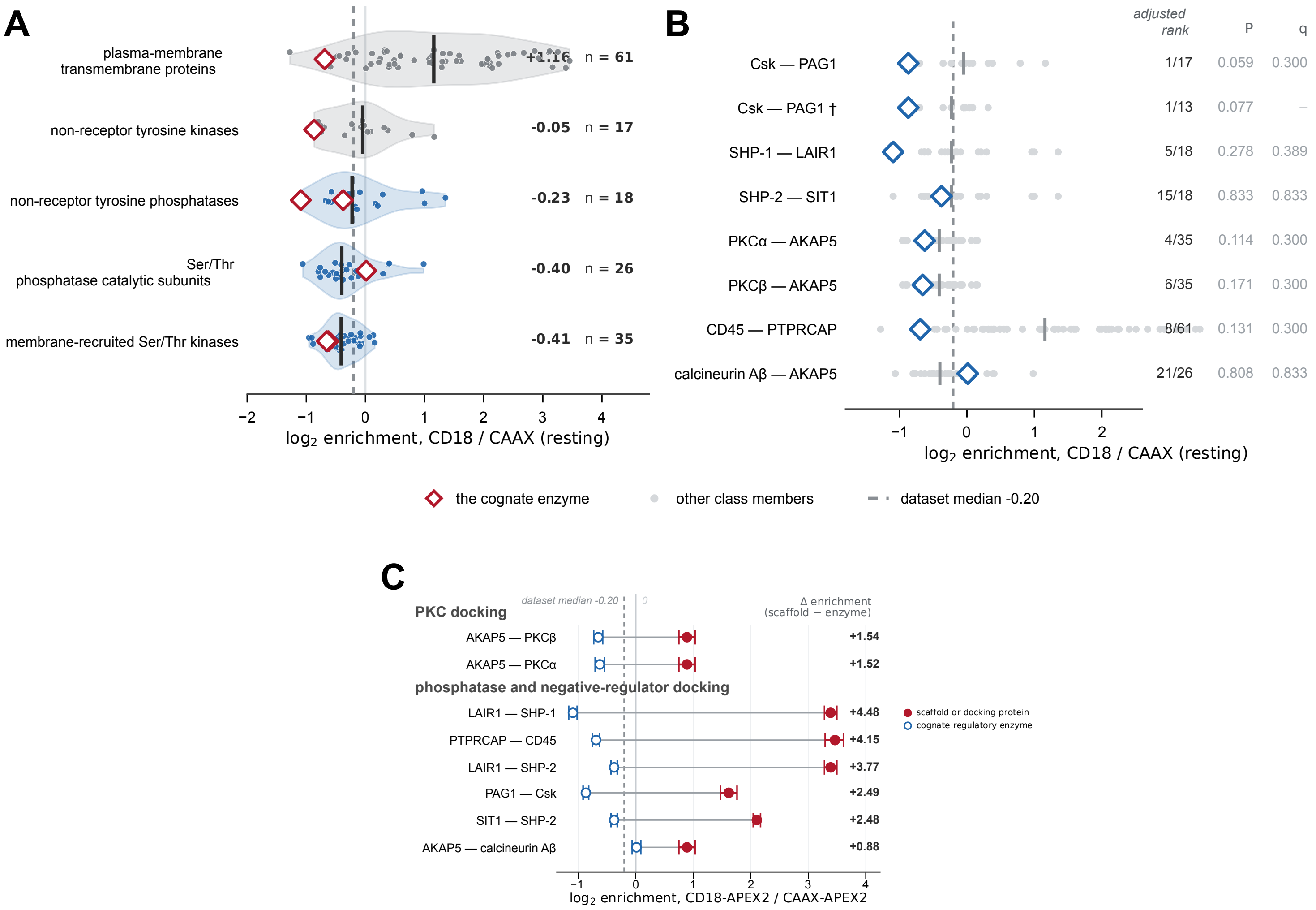

**Supplementary Fig. 4 | Enzyme proximity to the resting CD18 territory is dominated by protein-class architecture**

**a,** Resting CD18/CAAX enrichment distributions for plasma-membrane transmembrane proteins (n = 61) and four catalytic enzyme classes: non-receptor tyrosine kinases (n = 17), non-receptor tyrosine phosphatases (n = 18), Ser/Thr phosphatase catalytic subunits (n = 26) and membrane-recruited Ser/Thr kinases (n = 35). Points denote individual proteins, black bars denote class medians (+1.16, −0.05, −0.23, −0.40 and −0.41 $\log_2$, respectively) and the dashed line denotes the whole-dataset median (−0.20 $\log_2$). Red diamonds mark the cognate enzymes considered in b. **b,** Resting enrichment of the reported cognate enzymes relative to the other members of their class. Horizontal position shows CD18/CAAX enrichment; the adjusted rank is calculated within class after leave-one-out adjustment for captured abundance and peptide depth. Exact one-sided P values and Benjamini–Hochberg-adjusted q values are given for the seven primary enzyme tests. The daggered Csk–PAG1 entry is a sensitivity analysis excluding lipid-anchored Src-family kinases from the kinase class and was not included in the multiple-testing family. **c,** Resting enrichment of literature-defined docking proteins (red) and the regulatory enzymes they are reported to recruit (blue). Points show fitted $\log_2$ CD18-APEX2/CAAX-APEX2 enrichment with model-based uncertainty; values at right give the difference in enrichment between scaffold and enzyme. Connectors denote literature-defined relationships, not interactions measured by APEX2, and rows are descriptive examples rather than independent statistical pairs. Calcineurin Aβ (PPP3CB; 12 peptides, +0.01) is shown as the best-measured catalytic subunit; Aα (PPP3CA; 3 peptides, +0.98) and Aγ (PPP3CC; 4 peptides, −0.27) are given in the source data.

### Supplementary Figure 5

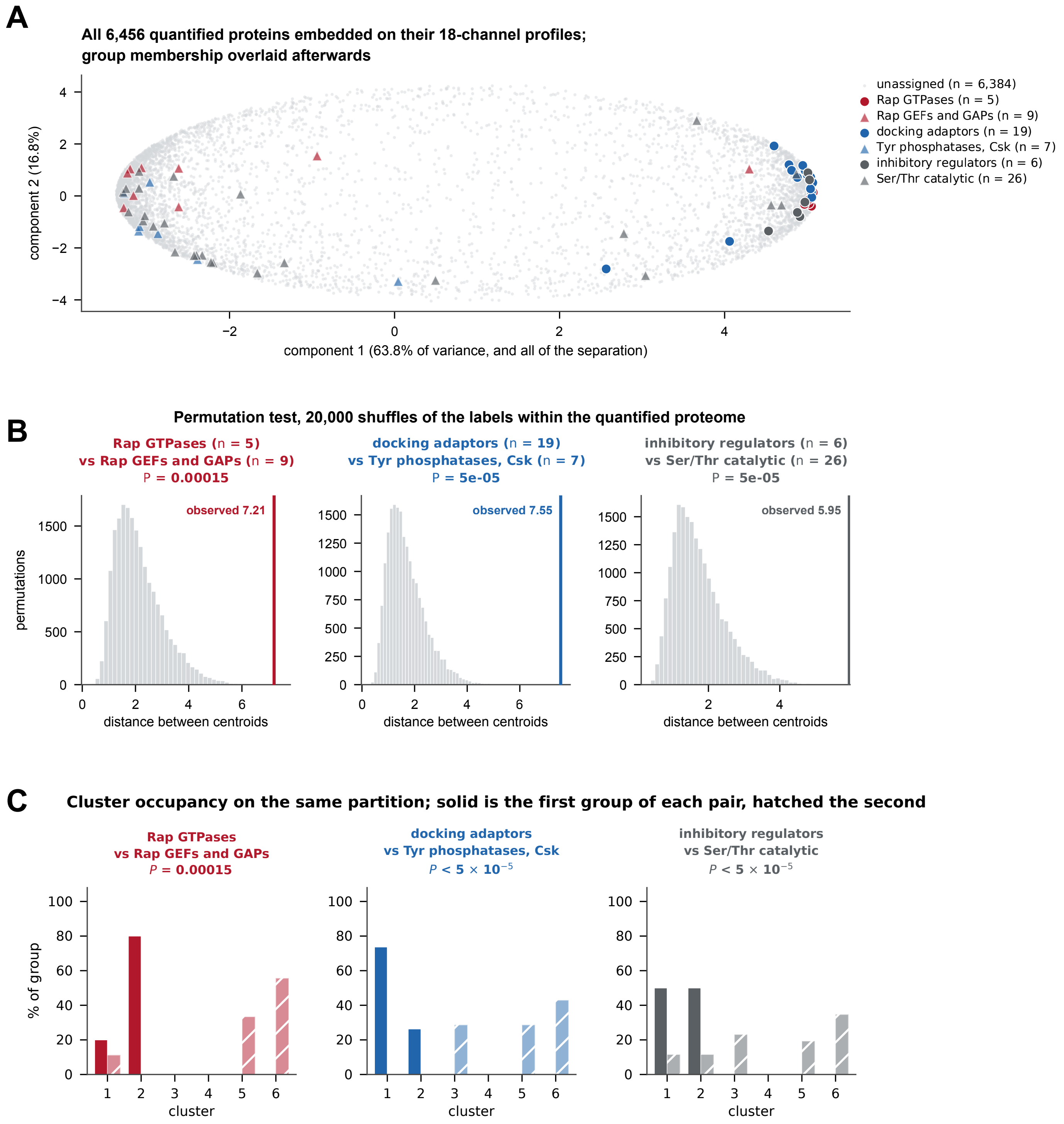

**Supplementary Fig. 5 | Functionally opposed regulatory classes occupy distinct regions of the CD18 proximity landscape**

**a,** Principal-component representation of all 6,456 quantified proteins using their complete 18-channel abundance profiles. Functional-class membership was overlaid after embedding: Rap GTPases (n = 5), Rap exchange factors and GAPs (n = 9), docking adaptors (n = 19), tyrosine phosphatases and Csk (n = 7), activity-restraining and regulatory proteins (n = 6) and Ser/Thr phosphatase catalytic subunits (n = 26); unassigned proteins (n = 6,384) are gray. Components 1 and 2 account for 63.8% and 16.8% of the variance, respectively, and all of the class separation lies along component 1. **b,** Empirical null distributions of the distance between class centroids from 20,000 random permutations of class labels within the quantified proteome. Vertical lines mark the observed distances for Rap GTPases versus Rap exchange factors and GAPs (7.21; $P = 1.5 \times 10^{-4}$), docking adaptors versus tyrosine phosphatases and Csk (7.55; $P \leq 5 \times 10^{-5}$) and activity-restraining and regulatory proteins versus Ser/Thr phosphatase catalytic subunits (5.95; $P \leq 5 \times 10^{-5}$). Exact permutation P values; $5 \times 10^{-5}$ is the resolution floor of 20,000 permutations. **c,** Occupancy of a six-cluster partition of the same embedding by each paired class. Solid bars denote the first and hatched bars the second class in each pair. n denotes proteins assigned to each class.

### Supplementary Figure 6

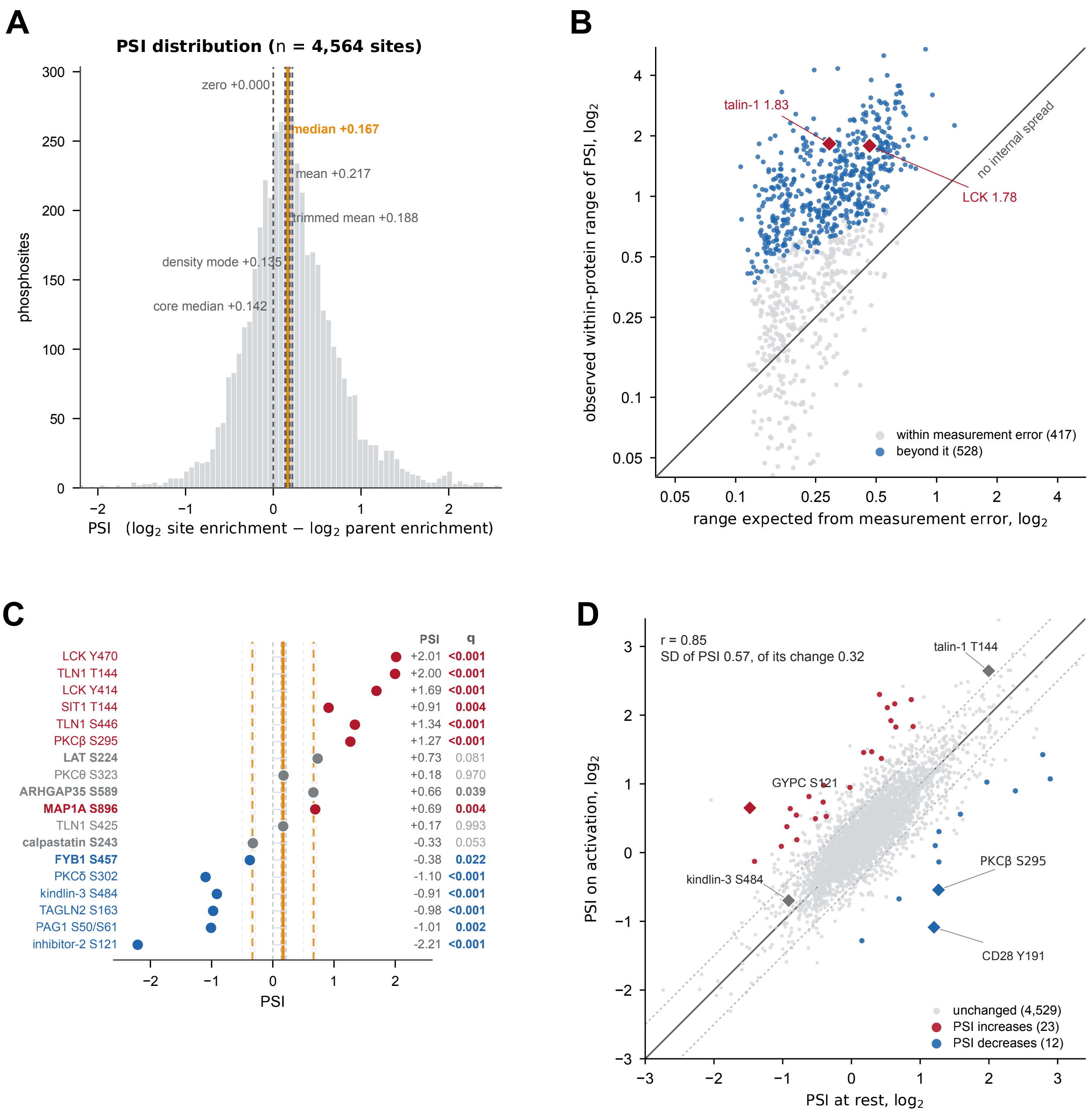

**Supplementary Fig. 6 | Parent-corrected phosphosite selectivity reveals within-protein heterogeneity that is largely preserved on activation**

**a,** Distribution of resting PSI across all 4,564 site-parent pairs. Vertical lines mark zero and alternative estimates of the distribution center: the median (+0.167, used throughout), mean (+0.217), trimmed mean (+0.188), density mode (+0.135) and core median (+0.142). **b,** Observed within-protein range of PSI compared with the range expected from the measurement uncertainty of the constituent sites, for the 945 proteins with more than one quantified phosphosite. Blue denotes proteins whose internal PSI spread exceeds measurement error (n = 528) and gray those that do not (n = 417); the diagonal denotes no internal spread beyond error. Talin-1 (1.83 $\log_2$) and LCK (1.78 $\log_2$) are highlighted. **c,** Resting PSI for selected phosphosites spanning receptor-preferential, parent-tracking and receptor-disfavored states. Vertical lines mark the empirical PSI center (orange) and the ±0.5 $\log_2$ effect-size criterion; red and blue denote sites significantly above or below the center and gray denotes sites not distinguishable from it. PSI values and q values are given at right. q values are Benjamini–Hochberg adjusted across the 4,564 PSI tests. The kindlin-3 measurement refers to S484. q values below 0.001 are shown as < 0.001. **d,** PSI at rest against PSI after ICAM-1+PMA for 4,564 site-parent pairs. Red and blue denote sites with significant PSI increases (n = 23) and decreases (n = 12), respectively; gray denotes unchanged sites (n = 4,529). The solid line is identity and the dotted lines mark ±0.5 $\log_2$. Pearson r = 0.85; the s.d. of PSI is 0.57 $\log_2$ and that of its activation-associated change is 0.32 $\log_2$. Selected sites are labeled.

### Supplementary Figure 7

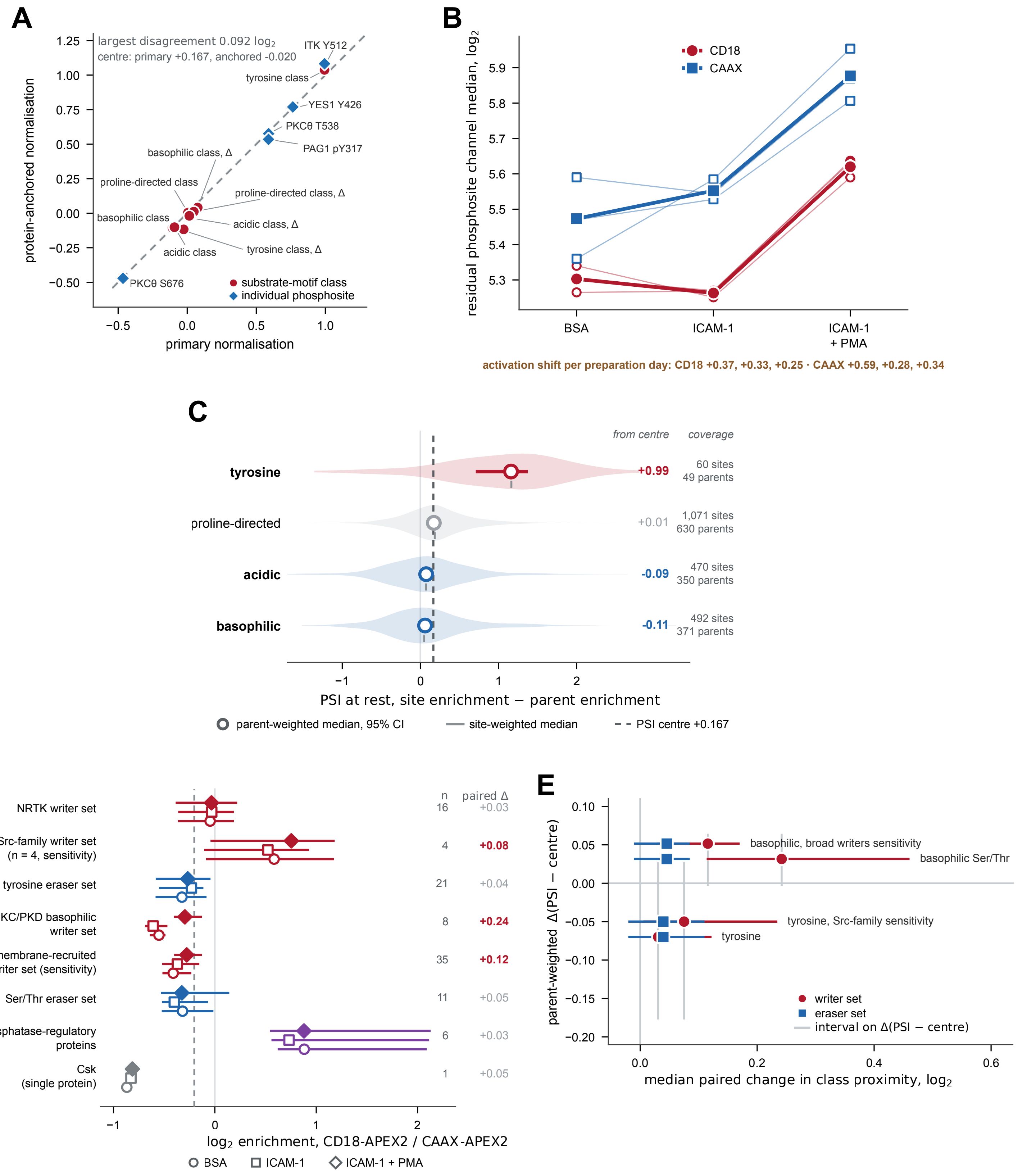

**Supplementary Fig. 7 | Writer and eraser proximity does not fully account for the phosphostate of the CD18 territory**

**a,** Comparison of PSI relative to its distribution center under the primary, independently normalized phosphosite workflow and a sensitivity analysis in which phosphosites were normalized with protein-derived channel factors. Red circles denote substrate-motif classes and their activation-associated changes (Δ) and blue diamonds selected individual phosphosites; the dashed line is identity. The resting PSI centers are +0.167 and −0.020 $\log_2$ under the two workflows and the largest absolute disagreement among the displayed quantities is 0.092 $\log_2$. **b,** Residual phosphosite-channel median after protein-derived loading correction across the three conditions. Open symbols denote the three independent preparation days and filled symbols the across-day trajectory for the CD18 (red) and CAAX (blue) samples. Activation-associated shifts for the individual days are +0.37, +0.33 and +0.25 $\log_2$ in CD18 and +0.59, +0.28 and +0.34 $\log_2$ in CAAX; independent phosphosite normalization removes this shared shift while preserving relative spatial phosphoform preferences. **c,** Resting PSI distributions for tyrosine, proline-directed, acidic and basophilic phosphosites. Violins show site-level distributions; open circles with horizontal intervals denote the parent-weighted median and its parent-bootstrap 95% confidence interval, with one value per parent protein; short bars denote site-weighted medians. The dashed line marks the empirical PSI center (+0.167 $\log_2$). Values at right give the parent-weighted deviation from the center and the site and parent coverage of each class. **d,** CD18-APEX2/CAAX-APEX2 enrichment of prespecified writer, eraser and phosphatase-regulatory protein sets across resting BSA (circles), ICAM-1 alone (squares) and ICAM-1+PMA (diamonds). Symbols denote set medians and horizontal intervals bootstrap 95% confidence intervals; Csk is shown separately as a single protein. The right-hand column gives the set size and the median paired per-protein change between resting and activated conditions; bold values have bootstrap confidence intervals excluding zero. These sets were defined for the substrate-class analysis and differ from the broader annotation classes in Supplementary Fig. 4. **e,** Median paired writer- or eraser-set redistribution between resting and activated conditions plotted against the corresponding parent-weighted change in the substrate class, Δ(PSI − center). The PSI center is condition-specific (+0.167 at rest and +0.212 after activation), and subtracting its movement isolates class-specific remodeling. Horizontal and vertical intervals are bootstrap 95% confidence intervals.

### Supplementary Figure 8

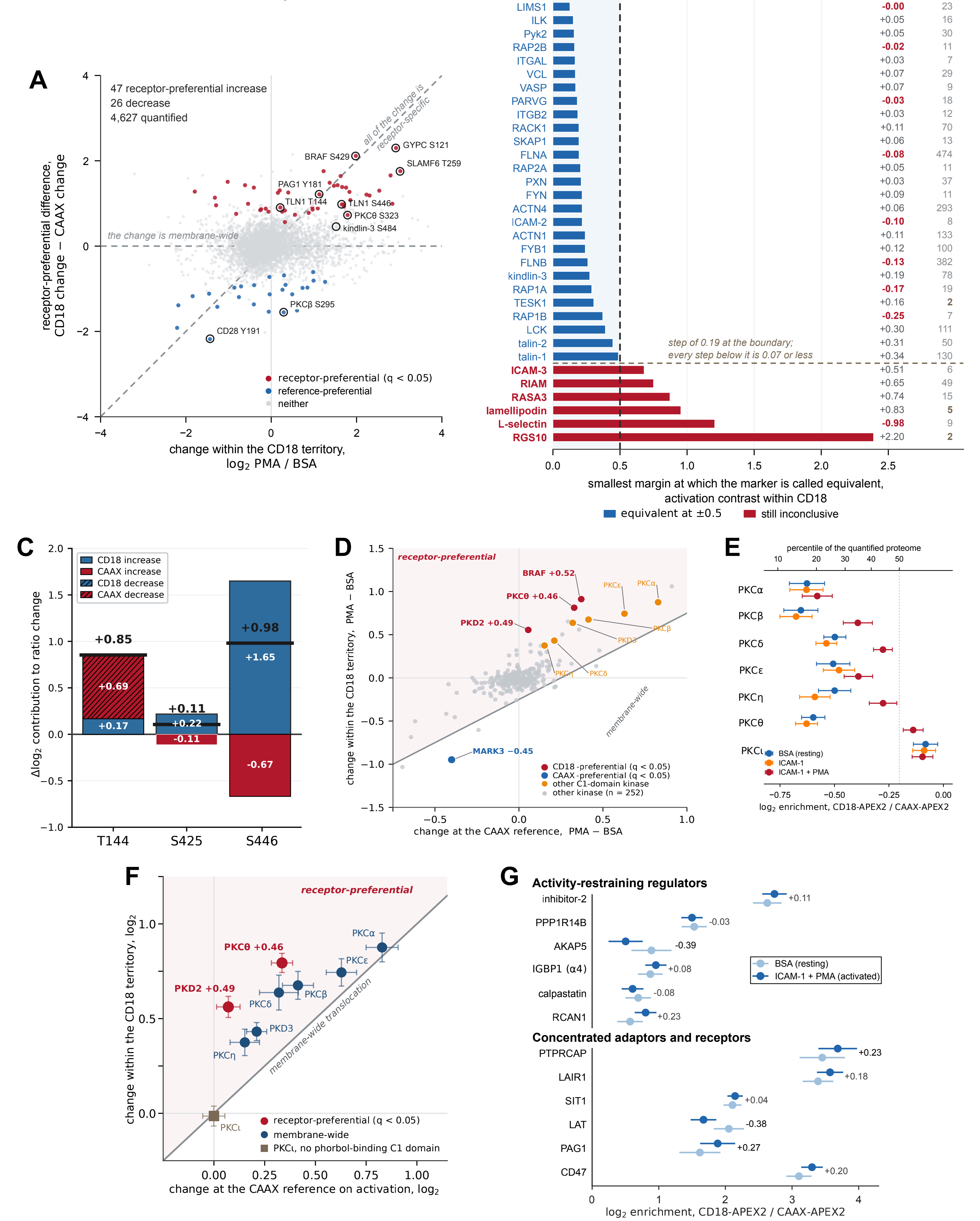

**Supplementary Fig. 8 | Activation remodeling shared with the CAAX reference, marker-set equivalence, kinase and PKC-family redistribution, and decomposition of the talin-1 phosphoform responses**

**a,** Receptor-preferential difference, defined as the CD18 change minus the corresponding CAAX change, plotted against the within-CD18 activation change for all 4,627 quantified phosphosites. Red and blue denote significant receptor-preferential increases (n = 47) and decreases (n = 26) at $q < 0.05$; gray denotes unresolved sites. The horizontal line corresponds to a response of equal magnitude in the CD18 samples and at the CAAX reference, and the diagonal to a response occurring entirely in the CD18 samples. Selected sites are labeled. **b,** Smallest symmetric equivalence margin at which each quantified canonical β2 integrin partner (34 of 36 quantified) is declared equivalent for the resting-versus-ICAM-1+PMA contrast within CD18. Blue denotes partners equivalent at the ±0.5 $\log_2$ margin used in this study (dashed line; n = 28) and red those that remain inconclusive at that margin (n = 6). Fitted activation changes and peptide counts are given at right. The six inconclusive partners form the upper end of a continuous ordering, separated from the remainder by a 0.19 $\log_2$ step at the boundary; every step below it is ≤ 0.07 $\log_2$. **c,** Decomposition of the resting-to-ICAM-1+PMA change in CD18/CAAX enrichment of the talin-1 T144, S425 and S446 phosphopeptides into the changes measured in the CD18 samples (blue) and at the CAAX reference (red). Because the receptor-preferential difference subtracts the CAAX change from the CD18 change, a decrease at the reference (hatched) contributes positively. Black horizontal marks and values above the bars give the net receptor-preferential difference (+0.85, +0.11 and +0.98 $\log_2$). For S446 the net difference combines a +1.65 $\log_2$ increase in the CD18 samples with a +0.67 $\log_2$ increase at the CAAX reference. **d,** Activation-associated change within the CD18 samples plotted against the corresponding change at the CAAX reference for all quantified protein kinases. Red and blue denote kinases with significant positive and negative receptor-preferential differences ($q < 0.05$; BRAF +0.52, PKD2 +0.49 and PKCθ +0.46 $\log_2$; MARK3 −0.45 $\log_2$), orange other C1 domain-containing kinases and gray the remaining kinases (n = 252). The diagonal corresponds to a change shared equally between the two labeling environments. **e,** CD18/CAAX enrichment of PKC-family proteins under resting BSA (blue), ICAM-1 alone (orange) and ICAM-1+PMA (red) conditions. The upper axis gives the corresponding percentile within the quantified proteome; the dashed line marks the dataset median. Points show fitted estimates with model-based uncertainty. Every phorbol-responsive isoform was under-represented at rest and increased on activation, whereas atypical PKCı, which lacks the phorbol-responsive C1-domain architecture, remained near parity. **f,** Activation-associated change within the CD18 samples plotted against the corresponding change at the CAAX reference for PKC and PKD family proteins. Red denotes isoforms with a significant receptor-preferential difference ($q < 0.05$; PKD2 +0.49 and PKCθ +0.46 $\log_2$); blue denotes isoforms whose change is shared between the two labeling environments. PKCı, which lacks a phorbol-binding C1 domain, is shown in brown. The diagonal corresponds to equal change in the two environments; the shaded region above it marks receptor-preferential responses. Points show fitted estimates with model-based uncertainty. **g,** Resting (light blue) and activated (dark blue) CD18/CAAX enrichment of the six selected activity-restraining and regulatory proteins and of six concentrated adaptors and receptors. Values at right give the activation-associated change in enrichment; none reached $q < 0.05$ (smallest $q = 0.33$, LAT). Measurements derive from three biological replicates.

### Supplementary Figure 9

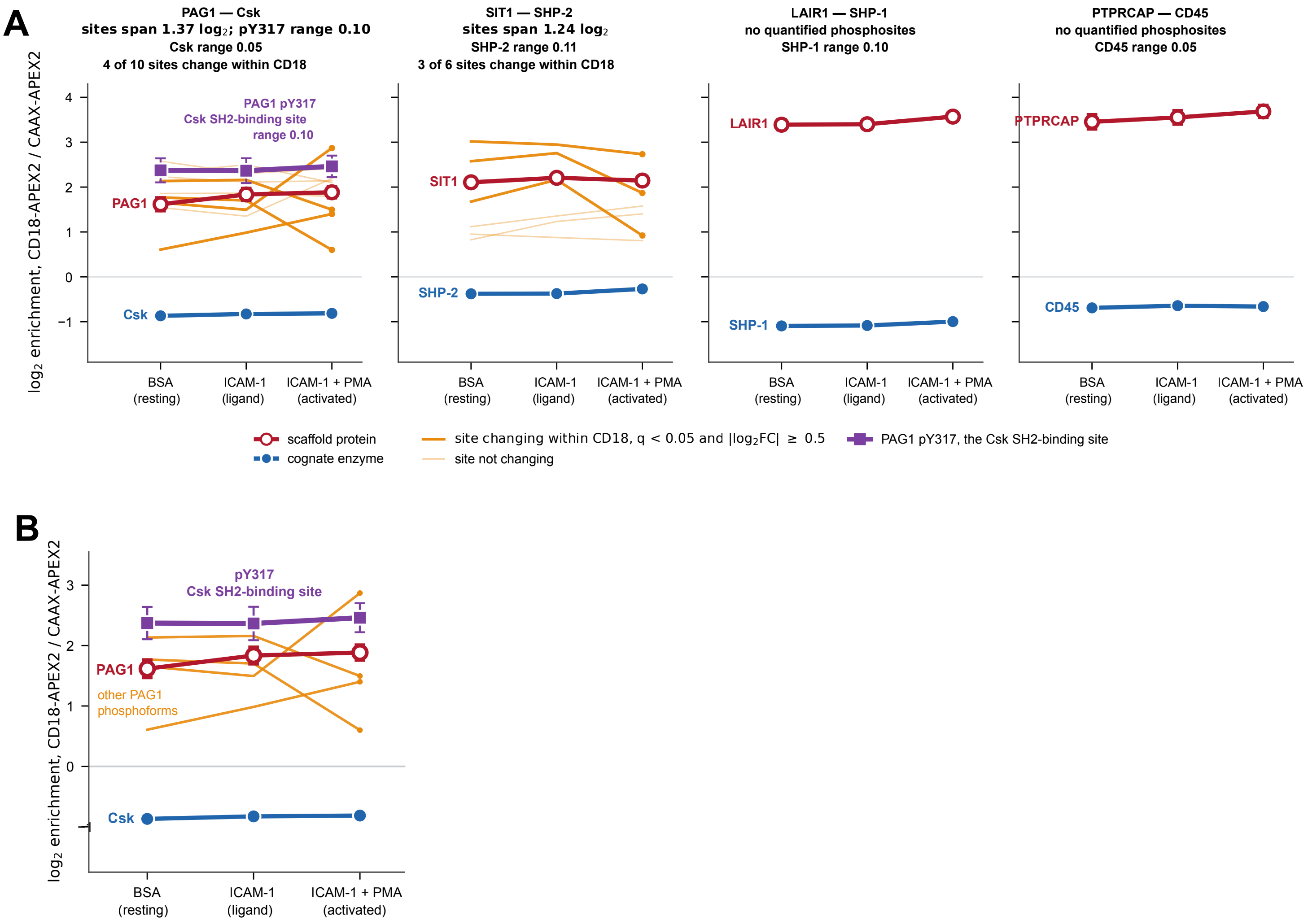

**Supplementary Fig. 9 | Scaffold phosphoform remodeling can occur without redistribution of the reported regulatory enzyme**

**a,** CD18/CAAX enrichment across resting BSA, ICAM-1 alone and ICAM-1+PMA conditions for the PAG1–Csk, SIT1–SHP-2, LAIR1–SHP-1 and PTPRCAP–CD45 pairs. Scaffold proteins are shown in red and the reported cognate enzymes in blue. Quantified scaffold phosphosites that change within the CD18 samples between resting and activated conditions ($q < 0.05$ and $|\log_2 \text{fold change}| \geq 0.5$) are shown as solid orange lines and non-changing sites as pale orange lines. The PAG1 phosphopeptide spanning the canonical Y317 Csk-docking region is shown in purple; the search localized the phosphate to Y317 (localization score 38.4), but S318 and S319 are adjacent alternative acceptors, so the residue assignment is provisional pending manual spectral confirmation. Individual PAG1 phosphosites vary by up to 1.37 $\log_2$ across conditions and four of ten change within CD18, whereas the Y317-region phosphopeptide and Csk vary by 0.10 and 0.05 $\log_2$, respectively. SIT1 phosphosites span 1.24 $\log_2$ and three of six change, whereas SHP-2 varies by 0.11 $\log_2$. No phosphosites passing quantification filters were recovered for LAIR1 or PTPRCAP; SHP-1 and CD45 vary by 0.10 and 0.05 $\log_2$. **b,** The PAG1–Csk pair from a at larger scale. Points show fitted estimates with model-based uncertainty. APEX2 reports time-integrated proximity and does not distinguish persistent under-representation from transient enzyme encounter.

### Supplementary Figure 10

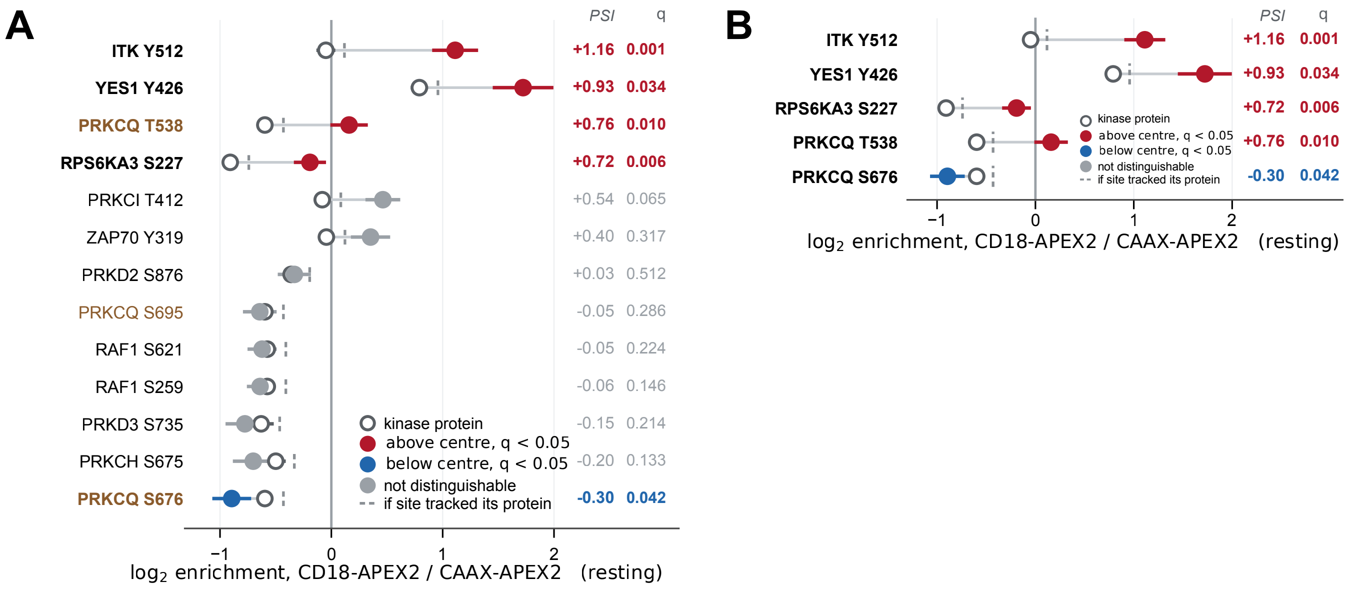

**Supplementary Fig. 10 | Selected kinase phosphoforms are spatially selected relative to their parent kinases**

**a,** Resting CD18-APEX2/CAAX-APEX2 enrichment of 13 prespecified kinase phosphosites (filled symbols) and their parent kinases (open circles). Short dashed ticks mark the position expected if each phosphosite tracked its parent protein at the empirical PSI center (+0.167 $\log_2$). Red and blue denote PSI significantly above or below the center, respectively; gray denotes no significant deviation. PSI and q values are given at right; q values are Benjamini–Hochberg adjusted across all 4,564 quantified site-parent pairs. The site set was defined before inspection of PSI values. ITK Y512 refers to the reference sequence used here and corresponds to the site commonly reported as Y511. The YES1 Y426 measurement derives from the Src-family activation-loop phosphopeptide LIEDNEpYTAR, whose sequence is shared among several family members, and is assigned to YES1 by parsimony. **b,** The five phosphosites from a that deviate significantly from their parent-relative expectation.
